# In Situ UNIversal Orthogonal Network (UNION) Bioink Deposition for Direct Delivery of Corneal Stromal Stem Cells to Corneal Wounds

**DOI:** 10.1101/2024.09.19.613997

**Authors:** Lucia G. Brunel, Betty Cai, Sarah M. Hull, Uiyoung Han, Thitima Wungcharoen, Gabriella Maria Fernandes-Cunha, Youngyoon Amy Seo, Patrik K. Johansson, Sarah C. Heilshorn, David Myung

## Abstract

The scarcity of human donor corneal graft tissue worldwide available for corneal transplantation necessitates the development of alternative therapeutic strategies for treating patients with corneal blindness. Corneal stromal stem cells (CSSCs) have the potential to address this global shortage by allowing a single donor cornea to treat multiple patients. To directly deliver CSSCs to corneal defects within an engineered biomatrix, we developed a UNIversal Orthogonal Network (UNION) collagen bioink that crosslinks *in situ* with a bioorthogonal, covalent chemistry. This cell-gel therapy is optically transparent, stable against contraction forces exerted by CSSCs, and permissive to the efficient growth of corneal epithelial cells. Furthermore, CSSCs remain viable within the UNION collagen gel precursor solution under standard storage and transportation conditions. This approach promoted corneal transparency and re-epithelialization in a rabbit anterior lamellar keratoplasty model, indicating that the UNION collagen bioink serves effectively as an *in situ*-forming, suture-free therapy for delivering CSSCs to corneal wounds.

**TEASER.** Corneal stem cells are delivered within chemically crosslinked collagen as a transparent, regenerative biomaterial therapy.

## INTRODUCTION

Over 12.5 million people globally suffer from corneal blindness, a leading cause of visual impairment.(*1*) Unlike cataracts or glaucoma which develop later in life, corneal blindness occurs after acute injury or disease and thus generally affects younger patients, with a devastating impact on their quality of life.(*2*) The only known curative treatment is corneal transplantation, yet the cadaveric human donor tissue required for this procedure is unfortunately available to less than 2% of patients worldwide.(*1*) To address this major global health need, we aimed to engineer corneal cell-laden collagen hydrogels as a curative treatment for corneal blindness that does not require a 1:1 donor cornea-to-patient ratio.

Recently, injection of corneal stromal stem cells (CSSCs) into the cornea has empirically demonstrated improved re-epithelialization of the corneal surface and reduced formation of scar tissue after injury.(*3*, *4*) CSSCs, also referred to as corneal mesenchymal stromal cells, are the progenitor cells for the corneal stroma.(*5*) They are derived from the corneal limbus and can differentiate into keratocytes, the quiescent stromal cells characteristic of healthy cornea.(*5*) Previous investigations of CSSCs have demonstrated that these cells can have antifibrotic effects, prevent the development of corneal scarring after injury, and help suppress inflammation, remodel pathological stromal tissue, and restore transparency.(*3*, *6*, *7*) Importantly, CSSCs are readily isolated and cultured from both donor and autologous sources and as such can expand the therapeutic potential of graft tissue from one beneficiary to many.(*3*, *6*, *8–10*) Prior approaches for the delivery of CSSCs to wounded corneas have involved direct injection into the corneal stroma(*3*) or application of the cells to the wound surface within a fibrin glue.(*6*, *7*) To date, however, methods of consistent and efficient delivery of CSSCs have not yet been established to allow for their clinical translation.

CSSCs are not yet in clinical use due to important challenges in their implementation, including the lack of a suitable carrier to deliver and retain them at the wound site while also encouraging their pro-regenerative effects. Since cell therapies commonly suffer from poor retention of cells at the delivery site, hydrogels may be used as cell carriers to provide a suitable microenvironment for cell survival and engraftment(*11*) and as matrices to fill corneal defects and reconstruct lost tissue after removal of a corneal scar.(*12–14*) For effective direct delivery of cells to the cornea, the hydrogel must meet several key criteria: It should be cytocompatible to support cell viability and function, maintain long-term transparency to aid in vision restoration, and exhibit structural stability to preserve the gel’s shape and size, even under cell-induced forces that could otherwise cause gel deformation and detachment from the host-graft interface.(*15*, *16*)

Furthermore, unlike cell-only or gel-only therapies, cell-gel therapies introduce additional requirements for the preparation of the cell-laden hydrogel.(*17*) The hydrogel precursor solution must be cytocompatible for the encapsulated cells to remain viable during storage and transportation conditions necessary for clinical translation,(*18*) and the gelation process within the cornea must be gentle and efficient to avoid harming the encapsulated cells or the surrounding host tissue.(*19*) Indeed, a major challenge with previous approaches to engineered corneal tissues is the cytotoxicity of many common techniques for matrix crosslinking. Strategies for matrix crosslinking in the cornea have often leveraged photocrosslinking (*e.g.*, free radical polymerization of methacrylate groups), which can have off-target effects from unintended side reactions.(*13*, *20–22*) While photochemistry is FDA-approved for use in certain clinical instances such as corneal crosslinking (CXL) for patients with corneal ectatic disorders, this photocrosslinking process has well-established cytotoxic effects on keratocytes and corneal nerves.(*23*, *24*) Caution is therefore merited when considering chemical interventions on pathologically wounded corneas.

As a cell-friendly, biomaterial crosslinking strategy, we recently developed a UNIversal Orthogonal Network (UNION) approach that leverages a strain-promoted azide-alkyne cycloaddition (SPAAC) reaction to crosslink cell-laden bioinks for 3D bioprinting.(*25*, *26*) SPAAC is a bioorthogonal form of click chemistry that exhibits no cross-reactivity with cells and proteins; does not require a catalyst, light energy, or heat for initiation; and produces no toxic byproducts.(*27*, *28*) These features make UNION bioinks ideally suited for clinical translation of implants with encapsulated cells. Previously, UNION bioinks were used to 3D bioprint cell-laden constructs *ex situ*.(*25*, *26*) However, for corneal trauma in which the damage occurs in a localized region and surrounding tissue remains healthy, it is advantageous to replace only the wounded tissue; gels that form *in situ* can conform exactly to the wound size and avoid extensive surgery and the use of sutures.(*12–14*) We have now adapted the UNION crosslinking technique for collagen bioinks—the primary protein component of the cornea—to enable the direct delivery of CSSCs via *in situ* gelation within corneal wounds.

Our work seeks to address the major clinical need for off-the-shelf therapeutic strategies to address corneal blindness by delivering regenerative CSSCs to corneal defects, without further injuring or inflaming an already wounded cornea. We hypothesized that UNION collagen bioinks could serve as a hydrogel carrier for CSSCs that gels *in situ* through bioorthogonal, covalent chemistry. Unlike conventional, physically self-assembled collagen (PHYS collagen), the crosslinked UNION collagen is highly optically transparent and stable against forces from encapsulated CSSCs that can cause undesired contraction of the gel. The bioorthogonal nature of the crosslinking chemistry allows CSSCs to maintain high cell viability within a UNION collagen precursor solution under typical storage and transportation conditions. Furthermore, upon injection into corneal wounds, the bioorthogonal crosslinking chemistry proceeds rapidly under physiological conditions without off-target effects on surrounding tissue. Through *in vitro* and *in vivo* evaluations, we demonstrate the clinical potential of UNION collagen bioinks as an *in situ*- forming, catalyst-free, and suture-free therapy that directly delivers CSSCs to corneal wounds, promoting epithelial wound healing and corneal regeneration.

## RESULTS

### Design and synthesis of UNION collagen for corneal regeneration

To effectively restore proper vision, bioengineered gels that deliver CSSCs to corneal wounds must restore uniform transparency to the wounded cornea with robust structural stability (**Figure 1**). Collagen type I is a promising material for corneal tissue engineering due to its critical role in the native human cornea. Collagen comprises 71% of the dry weight of the human cornea, governing the biomechanical properties of the corneal stroma and providing a supportive extracellular microenvironment for corneal cells.(*29–31*) Furthermore, the low immunogenicity of collagen, especially atelocollagen, reduces the risk of adverse reactions, making it a safe and effective material for developing regenerative therapies.(*32*) Collagen products have previously been approved by the FDA for a variety of medical and cosmetic uses (*e.g.*, dermal fillers).(*33*)

**Figure 1.**
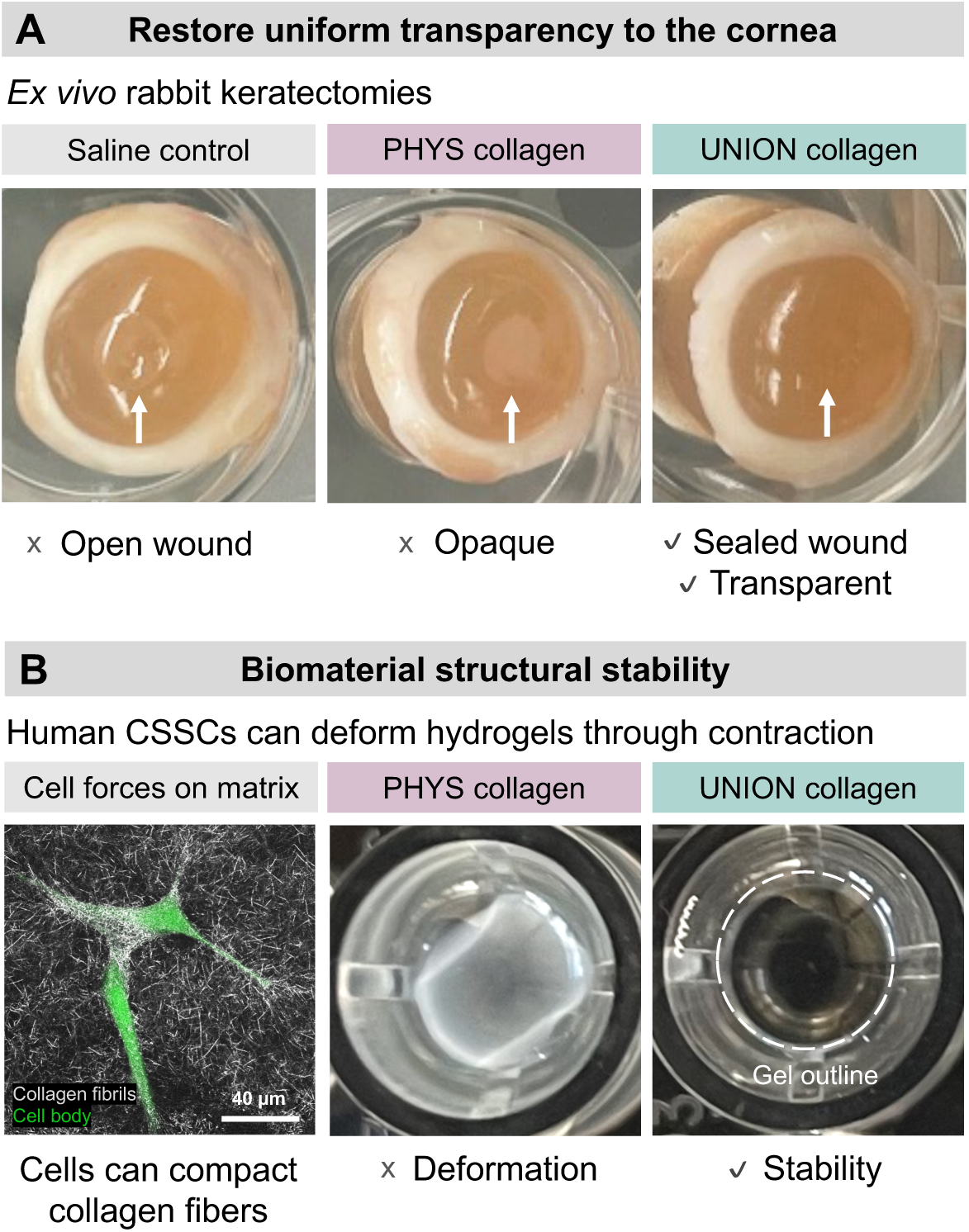
Key requirements for gels delivering corneal stromal stem cells (CSSCs) to corneal wounds. **(A)** The cell-gel therapy must seal the corneal wound and restore uniform transparency to the tissue. While both physically self-assembled (PHYS) and UNIversal Orthogonal Network (UNION) collagen bioinks can effectively fill corneal wounds, only UNION collagen provides an optically transparent matrix. **(B)** The cell-gel therapy must resist bulk deformation caused by CSSC-generated forces. PHYS collagen is prone to deformation, while UNION collagen remains highly stable.

Following a corneal injury, it is crucial to first seal the wound and restore transparency (**Figure 1A**). While conventional collagen that physically self-assembles into gels (PHYS collagen) can effectively fill an open corneal wound, these gels are characteristically opaque. Beyond demonstrating optical transparency, the gel must also maintain its structural stability, resisting deformation or contraction from cell-generated forces that could impair the gel’s engraftment at the implantation site (**Figure 1B**). Many cell types—including CSSCs—can exert significant tensions on their surrounding matrix.(*26*, *34–37*) When these cells are encapsulated within PHYS collagen, the gels are prone to significant deformation of their bulk shape and size.(*26*, *34–37*) Based on these results and previous *in vitro* material development studies,(*26*) we hypothesized that covalent, bioorthogonal crosslinking chemistry rather than physical self-assembly of collagen would allow us to meet key requirements of corneal tissue engineering, by filling corneal wounds with a stable and transparent gel that directly delivers regenerative CSSCs to the wound site.

Covalent crosslinking of hydrogels generally improves the material stability, helping maintain matrix properties over time. Since common hydrogel crosslinking strategies often exhibit cross-reactivity that can alter cell phenotype or even induce DNA damage or cell death, bioorthogonal chemistries have been recently developed as highly selective reactions that do not interfere with living cells.(*19*) We chose a bioorthogonal, covalent click chemistry that enables the UNION collagen bioinks to crosslink *in situ* to fill corneal wounds and directly deliver CSSCs on demand (**Figure 2**). Specifically, the UNION collagen is crosslinked with SPAAC chemistry between an azide and strained alkyne (**Figure 2A**). This copper-free click chemistry can proceed under physiological conditions and in the presence of cells. Azide groups were conjugated onto bovine type I atelocollagen using N-hydroxysuccinimide (NHS) chemistry to react with collagen’s primary amines. From a fluorescamine assay for unconjugated primary amines before and after the collagen-azide modification, the degree of functionalization was evaluated to be ∼60% (**Figure S1A**).

**Figure 2.**
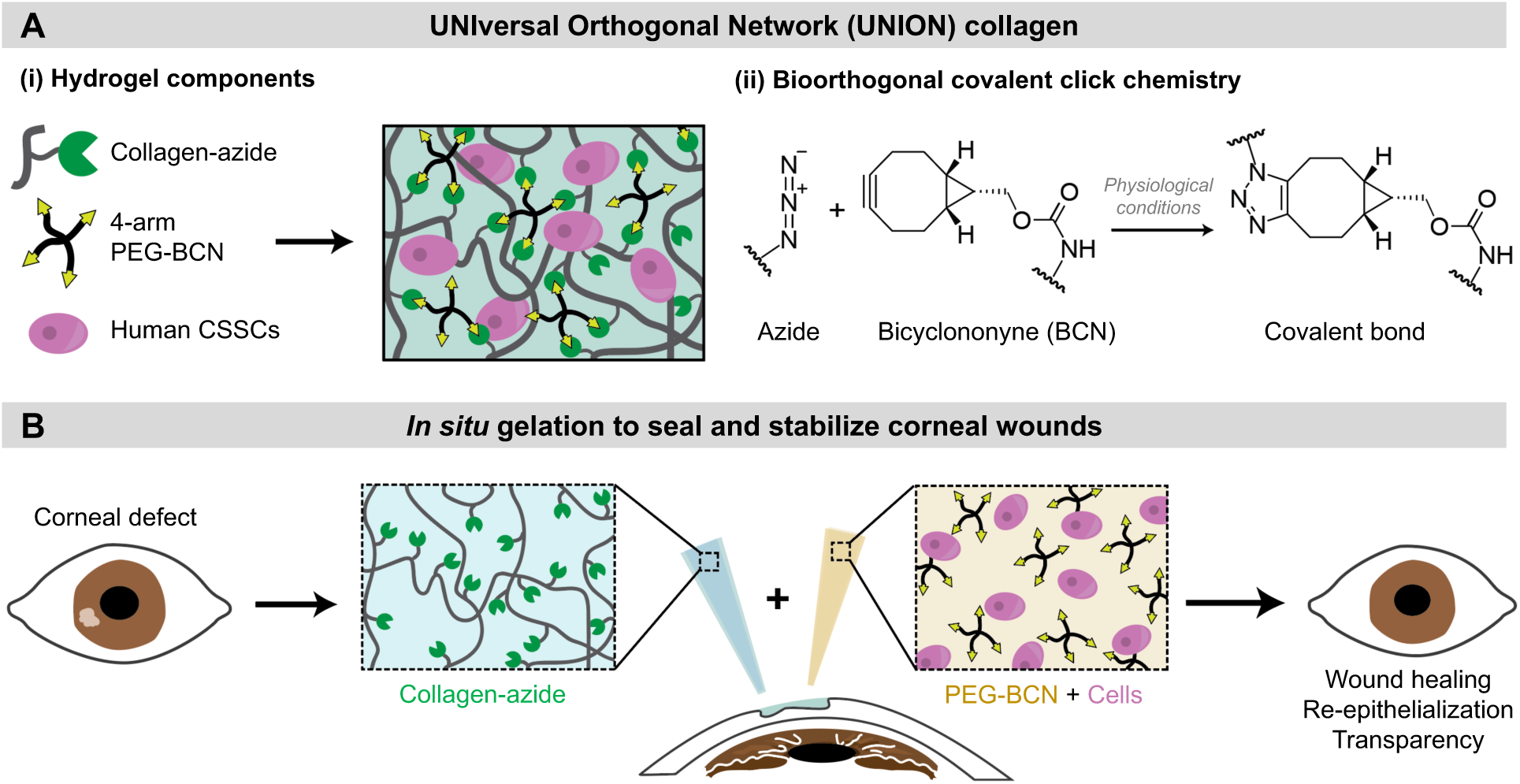
UNION collagen bioinks with encapsulated human CSSCs are designed to covalently gel *in situ* to promote corneal regeneration. (A) The UNION collagen bioink is composed of azide-modified collagen (collagen-azide), a 4-arm polyethylene glycol-bicyclononyne (PEG-BCN) crosslinker, and human CSSCs. The collagen-azide and PEG-BCN react under physiological conditions via strain-promoted azide- alkyne cycloaddition (SPAAC), a bioorthogonal click chemistry, to form covalent crosslinks. **(B)** The UNION collagen precursor components are mixed together and applied to the corneal defect to fill the wound with a CSSC-laden matrix.

To form the UNION collagen bioink, the collagen-azide is crosslinked with 4-arm polyethylene glycol-bicyclononyne (PEG-BCN), at concentrations of 3 mg/mL collagen-azide, 4 mg/mL PEG- BCN, and 3 million human CSSCs/mL. The UNION collagen bioink consists of two gel precursor solutions: (1) collagen-azide solution, and (2) PEG-BCN solution with human CSSCs. When mixed together and injected atop an open corneal wound, the UNION bioink material components undergo a crosslinking reaction *in situ* to create a 3D hydrogel matrix with the embedded CSSCs (**Figure 2B**). Due to the *in situ* gelation mechanism, the UNION collagen bioink conforms to the corneal wound size and shape, allowing for a suture-free surgery.

UNION collagen bioinks with encapsulated CSSCs have suitable mechanical and optical properties for corneal tissue engineering (**Figure 3**). The high cell viability of CSSCs pre-encapsulation is maintained during the encapsulation process and UNION collagen crosslinking (Day 0) as well as after 5 days of *in vitro* culture in UNION collagen (**Figure 3A**). In all cases, cell viability is > 95%, confirming the high cytocompatibility of the bioorthogonal SPAAC crosslinking and the resultant UNION collagen hydrogel. The UNION collagen gelates spontaneously upon mixing the collagen-azide and PEG-BCN crosslinker, reaching the crossover point (G’ = G”) in ∼10 minutes and a plateau storage modulus of ∼700 Pa (**Figure 3B**). The gelation kinetics and shear moduli are similar for UNION collage gels without cells (*i.e.* acellular) and those with encapsulated cells (*i.e.* cellular, 3 million CSSCs/mL), indicating that the presence of cells does not detectably affect or interfere with the hydrogel crosslinking.

**Figure 3.**
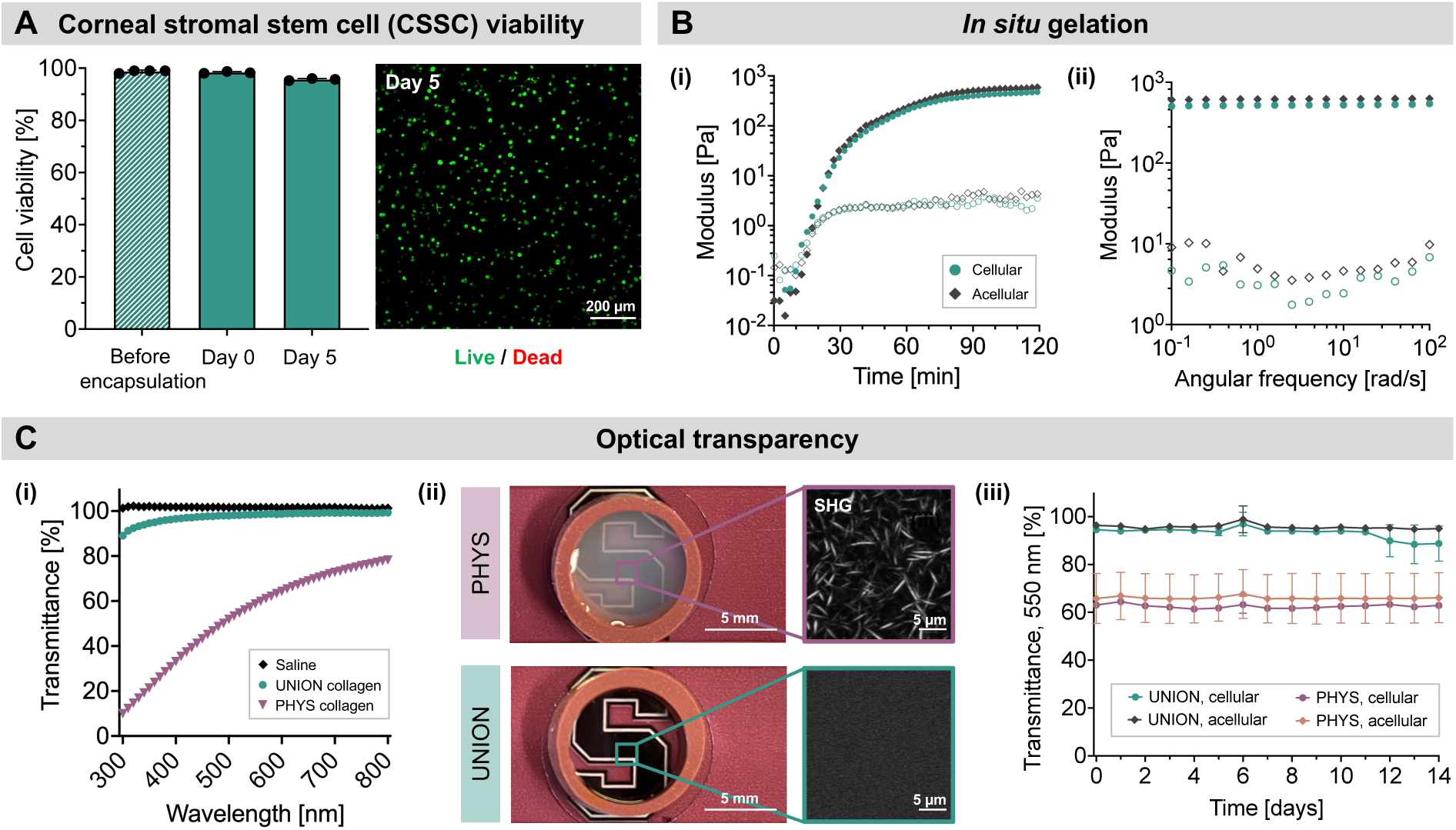
Cytocompatibility, gelation properties, and optical transparency of UNION collagen bioinks with encapsulated CSSCs. **(A)** CSSCs maintain their high cell viability after encapsulation and 5 days of *in vitro* culture within crosslinked, UNION collagen gels. N = 3 independent gels per condition. Data plotted as mean ± SD. **(B)** The presence of CSSCs does not interfere with the gelation of UNION collagen. (i) Representative gelation curves. (ii) Representative angular frequency sweeps. Filled symbols represent the storage modulus (G’), and open symbols represent the loss modulus (G”). **(C)** UNION collagen gels have high optical transparency. (i) Representative light transmittance curves in the visible light regime. (ii) Macroscopic transparency and microscopic structure of PHYS and UNION collagen gels. Second harmonic generation (SHG) imaging indicates the presence and lack of collagen fibrils for PHYS and UNION collagen, respectively. (iii) Cellular and acellular UNION collagen have higher light transmittance at 550 nm than cellular and acellular PHYS collagen across 14 days of *in vitro* culture. N = 3 independent gels per condition. Data plotted as mean ± SD.

Unlike conventional PHYS collagen gels, our UNION collagen gels exhibit the excellent optical transparency that is critical for corneal treatments (**Figure 3C**). As a control, PHYS collagen gels were prepared using the conventional technique of incubating neutralized bovine type I atelocollagen at 37 °C to allow a physical network of collagen fibrils to form. The PHYS collagen gels were formulated with 3 mg/mL collagen, the same concentration as the collagen-azide in the UNION collagen gels. While the light transmittance of PHYS collagen ranges between only 10- 80% transmittance across the wavelength range of visible light, the light transmittance of UNION collagen is above 90% transmittance for the same range of wavelengths. The stark improvement in transparency of UNION collagen is due to the difference in the microstructure of the gels, which was characterized with second harmonic generation (SHG) imaging of fibrils. The chemical modification and crosslinking of UNION collagen disrupts the physical self-assembly into fibrils,(*26*) thus preventing the challenges of light scattering from fibrils that causes PHYS collagen gels to be opaque.(*38*) The high transparency of UNION collagen compared to PHYS collagen is preserved across multiple concentrations of collagen (**Figure S1B**) and for both acellular and cellular (3 million CSSCs/mL) gels over 14 days in culture (Figure 3C, **Figure S1C**).

### *In vitro* characterization of UNION collagen with encapsulated corneal cells

Living cells dynamically interact with their surrounding hydrogel matrix, influencing both the structure and function of the material. This interaction is particularly critical in the context of corneal therapies, where the bulk size and geometry of the bioengineered gel affect its ability to fully fill the corneal wound and refract light properly. Mesenchymal stromal cells such as CSSCs are inherently contractile and can severely deform collagen hydrogels over time.(*26*, *34*) Therefore, the therapeutic potential of CSSCs within hydrogel carriers is currently limited in part by the undesired bulk deformation of gels in which CSSCs are delivered, since gel deformation can increase opacity and affect the gel interface with the native tissue. UNION collagen gels can overcome this challenge: While CSSCs rapidly and severely contract PHYS collagen gels, UNION collagen gels with the same density of cells over the same period of time in culture do not detectably contract (**Figure 4**).

**Figure 4.**
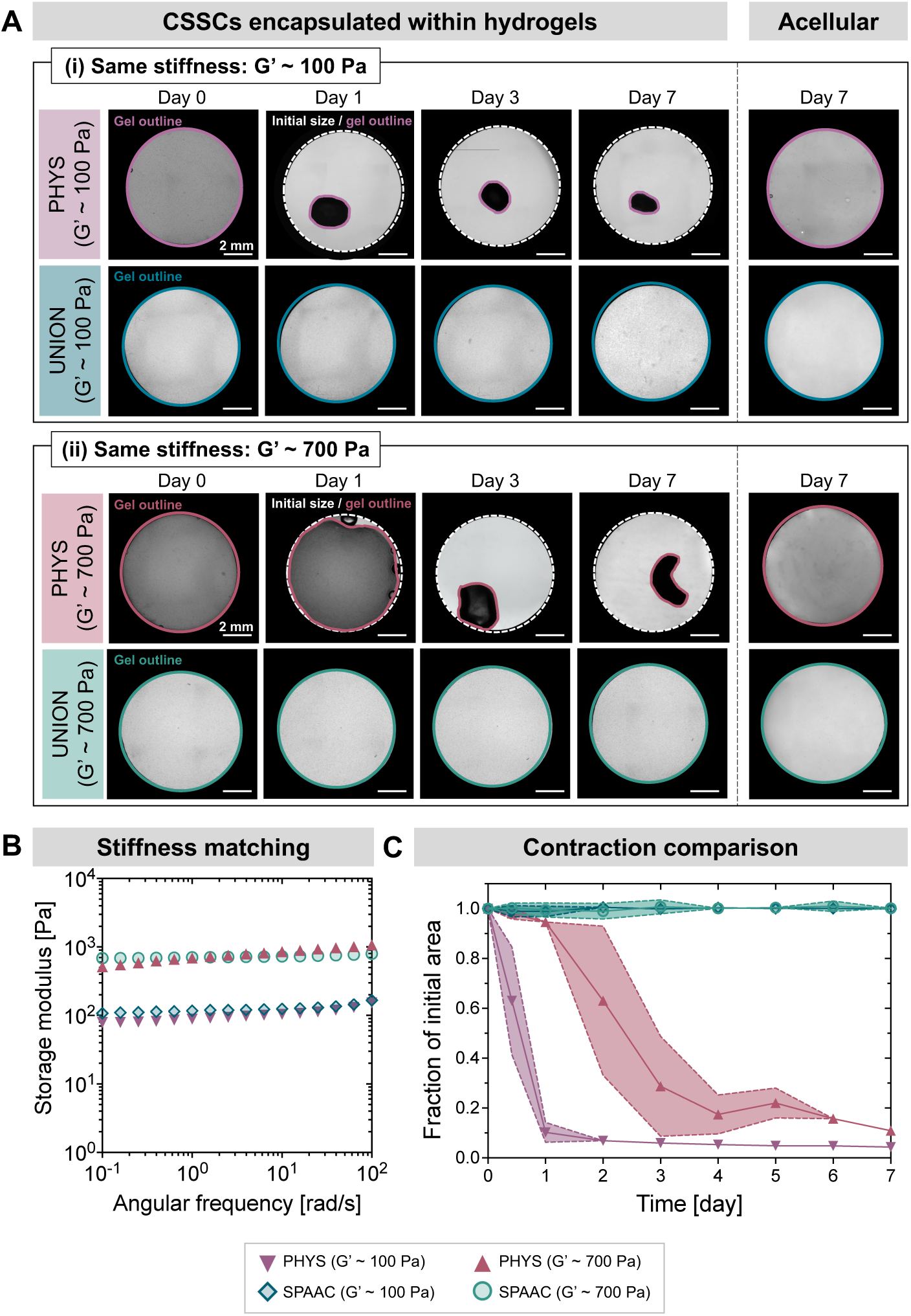
Cell-induced contraction of PHYS and UNION collagen hydrogels. **(A)** Representative images of the bulk size of CSSC-laden PHYS and UNION collagen gels (left) and acellular control gels (right) over time in culture. Scale bars = 2 mm. **(B)** Representative angular frequency sweeps of the PHYS and UNION collagen hydrogels, with formulations selected for stiffness-matched materials at ∼100 Pa and ∼700 Pa. PHYS collagen with G’ ∼ 100 Pa is 3 mg/mL collagen, PHYS collagen with G’ ∼ 700 Pa is 8 mg/mL collagen, UNION collagen with G’ ∼ 100 Pa is 3 mg/mL collagen-azide and 1.5 mg/mL PEG-BCN, and UNION collagen with G’ ∼ 700 Pa is 3 mg/mL collagen-azide and 4 mg/mL PEG-BCN. **(C)** Rate of collagen gel contraction. N ≥ 4 independent gels per material condition. Shaded regions represent the standard deviation from the mean.

Since PHYS collagen with 3 mg/mL collagen has a lower stiffness (G’ ∼ 100 Pa) than UNION collagen with 3 mg/mL collagen-azide (G’ ∼ 700 Pa), we formulated stiffness-matched conditions of PHYS and UNION collagen to isolate the effects of the crosslinking mechanism on cell-induced contraction. All cellular gels were prepared in circular 8-mm diameter molds with an initial concentration of 3 million CSSCs/mL, and the sizes of the gels were tracked daily over 7 days of culture (**Figure 4A**). To prepare gels with a stiffness of ∼100 Pa, PHYS collagen was formulated to be 3 mg/mL collagen, and UNION collagen was formulated to be 3 mg/mL collagen-azide with 1.5 mg/mL PEG-BCN. To prepare gels with a stiffness of ∼700 Pa, PHYS collagen was formulated to be 8 mg/mL collagen, and UNION collagen was formulated to be 3 mg/mL collagen-azide with 4 mg/mL PEG-BCN (**Figure 4B**).

Across both stiffness-matched pairs, the PHYS collagen gels suffered from severe contraction in size, while UNION collagen gels did not detectably contract (**Figure 4C**). On average, PHYS collagen gels with G’ ∼ 100 Pa contracted to ∼10% of their initial area within a single day and <5% of their initial area within 5 days. The PHYS collagen gels with higher stiffness (G’ ∼ 700 Pa) had slower contraction compared to PHYS collagen gels with lower stiffness (G’ ∼ 100 Pa), which is consistent with previous studies indicating that cell contractility is reduced in stiffer matrices.(*39*) However, PHYS collagen gels with G’ ∼ 700 Pa also experienced severe changes in size, contracting to < 30 % of their initial size within 3 days and ∼10% of their initial size after 7 days. The lack of any detectable contraction of UNION collagen gels observed at both stiffnesses (100 Pa and 700 Pa) indicates the robust stability against cell-induced contraction imparted by the covalent chemical crosslinks. Therefore, UNION collagen is well-suited as a hydrogel platform to deliver CSSCs and provide a microenvironment that remains structurally stable over time. Acellular gels did not change size for any material condition (Figure 4A), confirming that the contraction observed in cellular PHYS collagen gels is indeed caused by CSSC-generated forces.

To evaluate the regenerative potential of the cell-gel therapy, we assessed the phenotype of CSSCs encapsulated within the UNION collagen bioink and the growth of corneal epithelial cells over the gel’s surface (**Figure 5**). Immunocytochemistry analysis of CSSCs cultured within UNION collagen for 5 days confirmed the expression of aldehyde dehydrogenase 3A1 (ALDH-3A1), a corneal crystallin essential for maintaining cellular transparency and a characteristic marker of healthy CSSCs in the cornea (**Figure 5A**).(*40*) Additionally, the CSSCs produced hyaluronic acid, a native glycosaminoglycan that helps regulate corneal wound healing (**Figure 5B**).(*41*)

**Figure 5.**
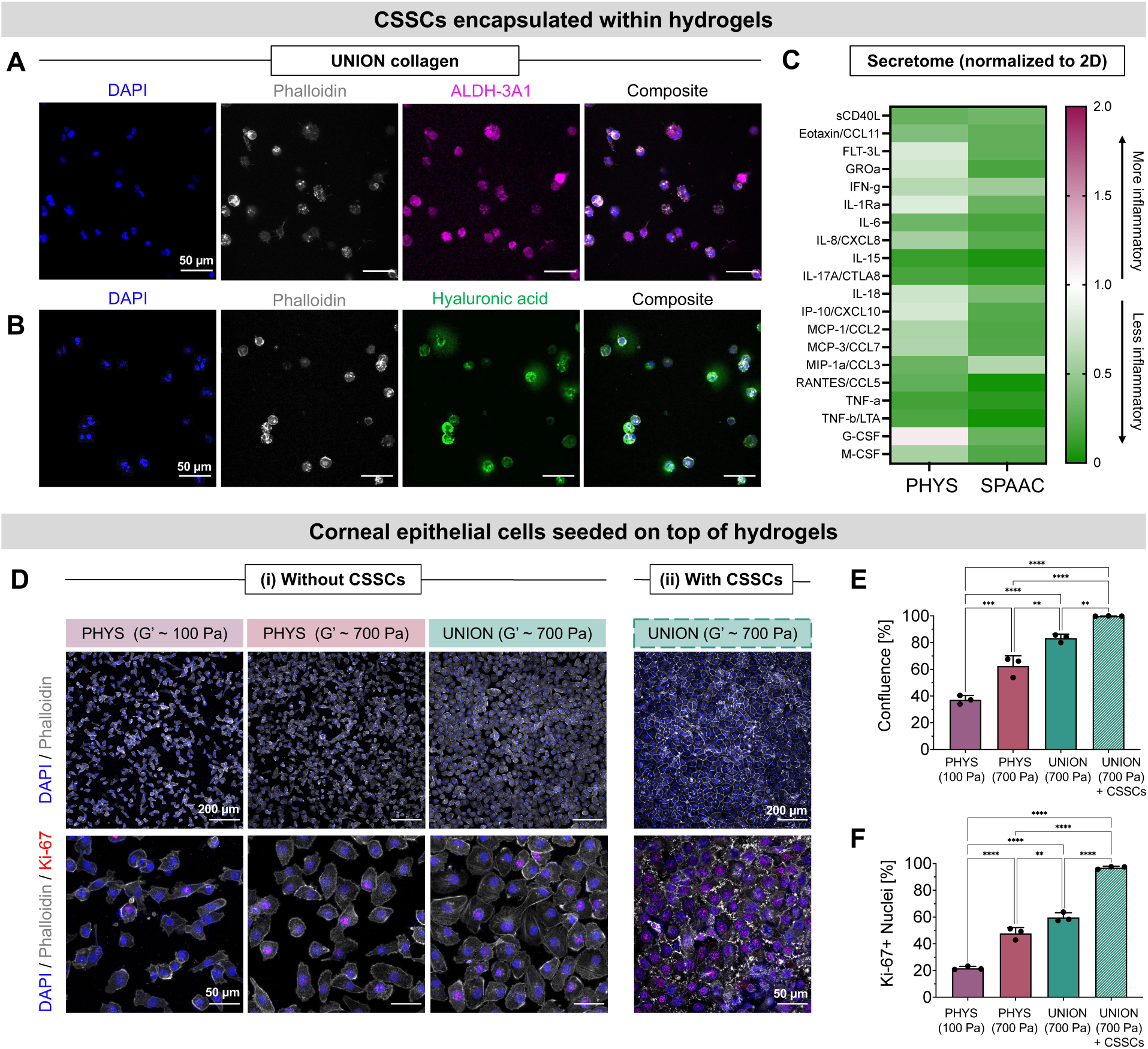
UNION collagen supports the encapsulation of CSSCs within the gel and the monolayer growth of corneal epithelial cells over the gel. **(A)** Staining of CSSCs encapsulated within UNION collagen for 5 days with DAPI (nuclei, blue), phalloidin (F-actin, gray), and anti-ALDH-3A1 (CSSC marker, magenta). Scale bars = 50 µm. **(B)** Staining of CSSCs encapsulated in UNION collagen for 5 days with DAPI (nuclei, blue), phalloidin (F-actin, gray), and biotinylated hyaluronic acid binding protein (hyaluronic acid, green). Scale bars = 50 µm. **(C)** Immunoassay of CSSC secretome for human cytokines, 5 days after CSSC encapsulation in PHYS and UNION collagen. Results are normalized to the same number of CSSCs on 2D tissue culture plastic. **(D)** Representative images of human corneal epithelial cells 2 days after seeding on PHYS and UNION collagen without (left) or with (right) encapsulated CSSCs. Staining with DAPI (nuclei, blue), phalloidin (F-actin, gray), and anti-Ki-67 (proliferation marker, red). Scale bars (top row) = 200 µm. Scale bars (bottom row) = 50 µm. **(E)** Quantification of corneal epithelial cell confluence. **(F)** Quantification of corneal epithelial cells positive for Ki-67. N = 3 independent gels per material condition. Statistical analyses were performed with an ordinary one-way ANOVA with Tukey’s multiple comparisons test. Data plotted as mean ± SD. \*\**p* < 0.01, \*\*\**p* < 0.001, \*\*\*\**p* < 0.0001.

It is known that corneal stromal cells cultured on 2D tissue culture plastic may undergo an undesired transition to a myofibroblast-like phenotype.(*42–44*) We therefore analyzed the CSSC secretome for the expression of pro-inflammatory cytokines (**Figure 5C**). Encapsulation of the CSSCs in 3D collagen matrices—both PHYS and UNION collagen—resulted in reduced secretion of pro-inflammatory cytokines compared to CSSCs cultured on 2D tissue culture plastic. This is consistent with previous work comparing the impact of 2D and 3D culture conditions on the therapeutic effects of mesenchymal stromal cell secretome for corneal wound healing.(*42*) Notably, CSSCs encapsulated in UNION collagen secreted even fewer pro-inflammatory cytokines than CSSCs in PHYS collagen. These findings highlight the advantages of UNION collagen matrices and their bioorthogonal crosslinking chemistry for preserving the regenerative phenotype of CSSCs while minimizing pro-inflammatory responses.

The proper regrowth of the corneal epithelium—the outermost layer of the cornea that serves as a barrier against pathogens, debris, and other harmful substances—is a crucial step in corneal regeneration.(*45*) We first evaluated the proliferation and spreading of human corneal epithelial cells seeded on top of acellular PHYS and UNION collagen gels (**Figure 5D**). Phalloidin staining of F-actin showed that the corneal epithelial cells spread evenly and confluently over the surface of UNION collagen. Immunocytochemistry further confirmed that the corneal epithelial cells on UNION collagen express cytokeratin 12 (CK-12), a characteristic marker of corneal epithelial cells that indicates differentiation into a mature epithelial phenotype (**Figure S2**).(*46*) We used a confluency assay to quantify the coverage of the corneal epithelial cells over the gels on Day 2 after seeding. The confluency on UNION collagen (UNION, G’ ∼ 700 Pa, confluency ∼ 80%) was significantly greater than that on (1) the PHYS collagen with the same concentration of 3 mg/mL collagen (PHYS, G’ ∼ 100 Pa, confluency ∼ 30%) and (2) the PHYS collagen with the higher concentration of 8 mg/mL collagen to reach the same stiffness as the UNION collagen (PHYS, G’ ∼ 700 Pa, confluency ∼ 50%) (**Figure 5E**). Therefore, even when the stiffnesses of the PHYS and UNION gels are the same, greater corneal epithelial cell growth is observed over the collagen gels that are covalently crosslinked (UNION) rather than physically assembled (PHYS), underscoring the utility of UNION crosslinking for gels promoting corneal re-epithelization.

CSSCs are known to exert some of their pro-regenerative effects in the cornea through secreted factors that allow crosstalk specifically with the corneal epithelial cells, thus promoting corneal re-epithelialization and wound healing.(*47*) Therefore, we next tested a co-culture system, with CSSCs encapsulated within the UNION collagen gel upon which the corneal epithelial cells grew. With the presence of CSSCs in the UNION collagen, the corneal epithelial cells were able to achieve confluency (∼98%) and adopt their characteristic cobblestone morphology even more effectively than over acellular UNION collagen gels (Figure 5D-E). These results suggested that the corneal epithelial cells may be most proliferative on the UNION collagen gels with encapsulated CSSCs, which we explored using Ki-67 immunostaining. Consistent with this hypothesis, we found that the proliferation of corneal epithelial cells was greater on UNION collagen than PHYS collagen, and on CSSC-laden UNION collagen than acellular UNION collagen (**Figure 5F)**. This demonstrates that both the UNION crosslinking technique and the delivery of CSSCs within the gel assist the spreading and proliferation of human corneal epithelial cells.

### Corneal cell viability in UNION collagen under shipping and storage conditions

The practical worldwide implementation of regenerative therapies with living cells requires standardized protocols for their storage and transportation (**Figure 6**). For conventional corneal transplants, corneal graft tissue is retrieved from cadaveric donors, stored at an eye bank, and then shipped overnight to surgery centers at which corneal operations are performed (**Figure 6A**). The most common storage method for corneal tissue in the United States of America and Asia is hypothermic storage, in which the tissue is maintained between 2-6 °C to reduce the metabolic activity of the cells.(*48*) Here, we demonstrate that the same infrastructure already in place for the shipping of donor corneal graft tissues would be suitable for the distribution of the UNION collagen bioink precursors with living CSSCs to surgery centers.

**Figure 6.**
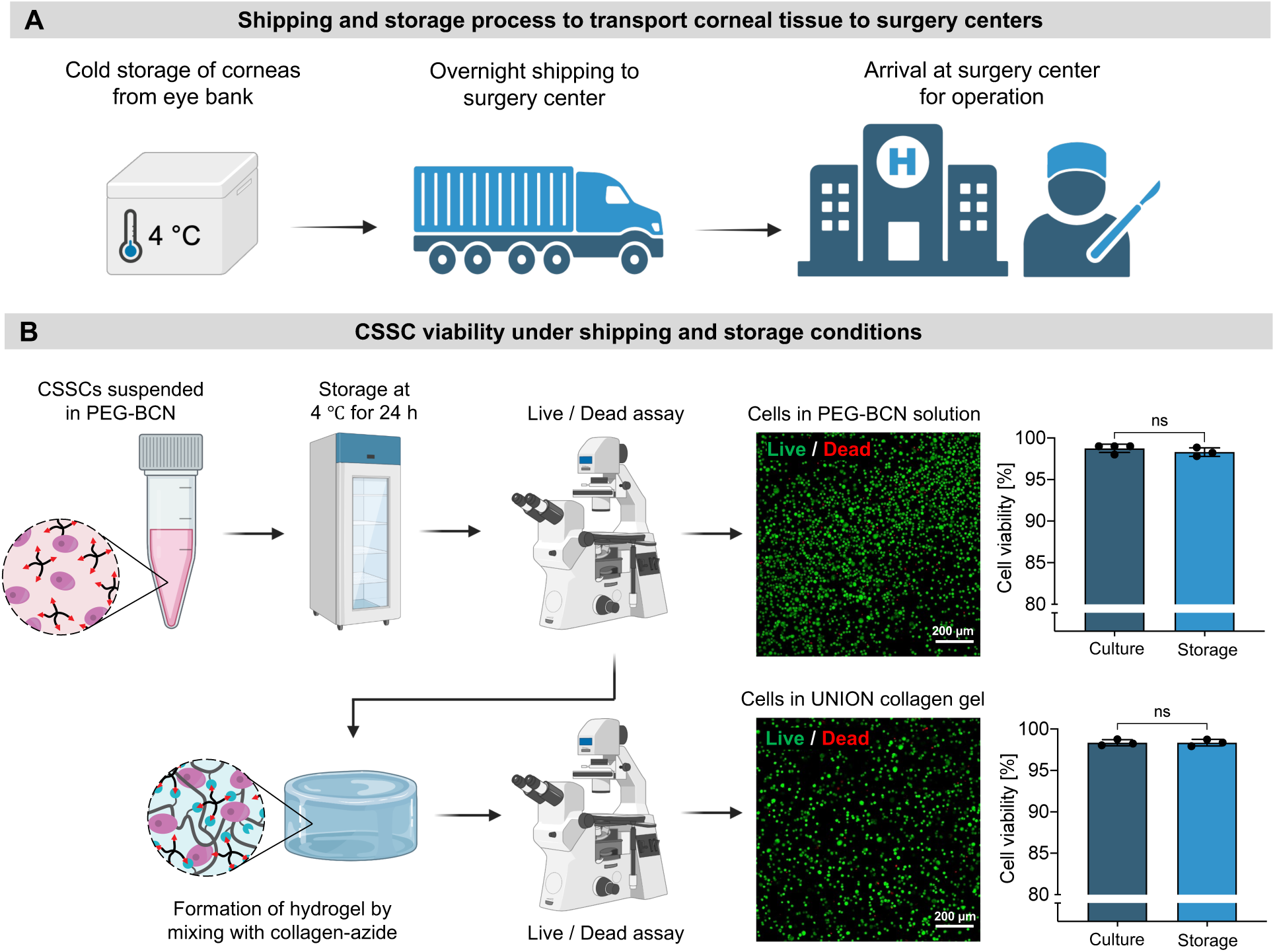
CSSCs remain viable under shipping and storage conditions. **(A)** Schematic illustrating the current storage and transportation process for donor corneas. Corneal tissues are stored at 4 °C and shipped overnight from eye banks to surgery centers for corneal transplantation surgeries. **(B)** After 24 h of cold storage at 4 °C, CSSCs remain highly viable both within the PEG-BCN solution (top) and after formation of the UNION collagen hydrogel by mixing with collagen-azide (bottom). N = 3 independent samples per condition. Normality of the data was confirmed with the Shapiro-Wilk test, and statistical analysis was performed with an unpaired t test. Data plotted as mean ± SD. ns = not significant.

To mimic shipping and storage conditions, the solution of collagen-azide was stored in one vial, while the solution of PEG-BCN mixed with CSSCs in cell culture medium was stored in a second vial. Both vials were kept at 4 °C for 24 hours to mimic overnight shipping on ice (**Figure 6B**). Following the 24 h cold storage, the CSSCs were evaluated with a Live / Dead cytotoxicity assay. Cells in the PEG-BCN solution immediately after 24-h cold storage were more than 95% viable, suggesting that high cell viability could be expected after shipping to a surgery center. Furthermore, once the vial of collagen-azide and the vial of PEG-BCN with CSSCs were mixed together and cast *in situ* to form a gel, the CSSCs continued to be more than 95% viable. This indicates that high cell viability could be expected after the cells are applied to corneal wounds at a surgery center within the UNION collagen bioink. In both cases—before and after UNION collagen gel formation—the viability of CSSCs subjected to 24 hours of cold storage was similarly high to control CSSCs prepared under normal culture conditions.

We then cultured the CSSCs that had undergone cold storage to assess their viability and phenotype at a later timepoint within the UNION collagen gel. After 5 days, the CSSCs within the UNION collagen gel maintained high cell viability (>95%) and expression of ALDH-3A1 (**Figure S3**). Furthermore, when the cold storage time was increased to 48 h or 72 h, mimicking shipping delays, CSSC viability after the extended cold storage time also remained greater than 95% (**Figure S4**). Therefore, the UNION collagen bioink system supports the high viability of CSSCs during standard storage and transportation conditions, including accounting for less-than-ideal shipping circumstances. The robust viability and stability of CSSCs within the UNION collagen bioink precursor solution under cold storage conditions support its potential for widespread clinical adoption. This offers a reliable and adaptable solution for distributing the cell-gel therapy that utilizes existing corneal transportation infrastructures.

### Validation of corneal regeneration in a rabbit anterior lamellar keratoplasty model

To evaluate the clinical potential of UNION collagen gels with CSSCs to promote corneal wound healing, we used a rabbit anterior lamellar keratoplasty model (**Figure 7, Figure S5**). No abnormalities were observed within the cornea or on the ocular surface prior to the operation (**Figure S5A**). An anterior keratectomy was performed to create a circular corneal wound with a diameter of 3.5 mm and a thickness of ∼200 µm (**Figure S5B**). This wound cut depth was controlled to be ∼50% of the thickness of the rabbit cornea (**Figure S5C**). For corneas treated with the UNION collagen gels with CSSCs, the solution of collagen-azide was mixed with the solution of PEG-BCN and CSSCs, and 5-7 µL of the resultant bioink were deposited within the corneal wound (**Figure S5D**). Since the treatment is applied as a solution, the bioink flows to fill the corneal wound size before crosslinking into a gel.

**Figure 7.**
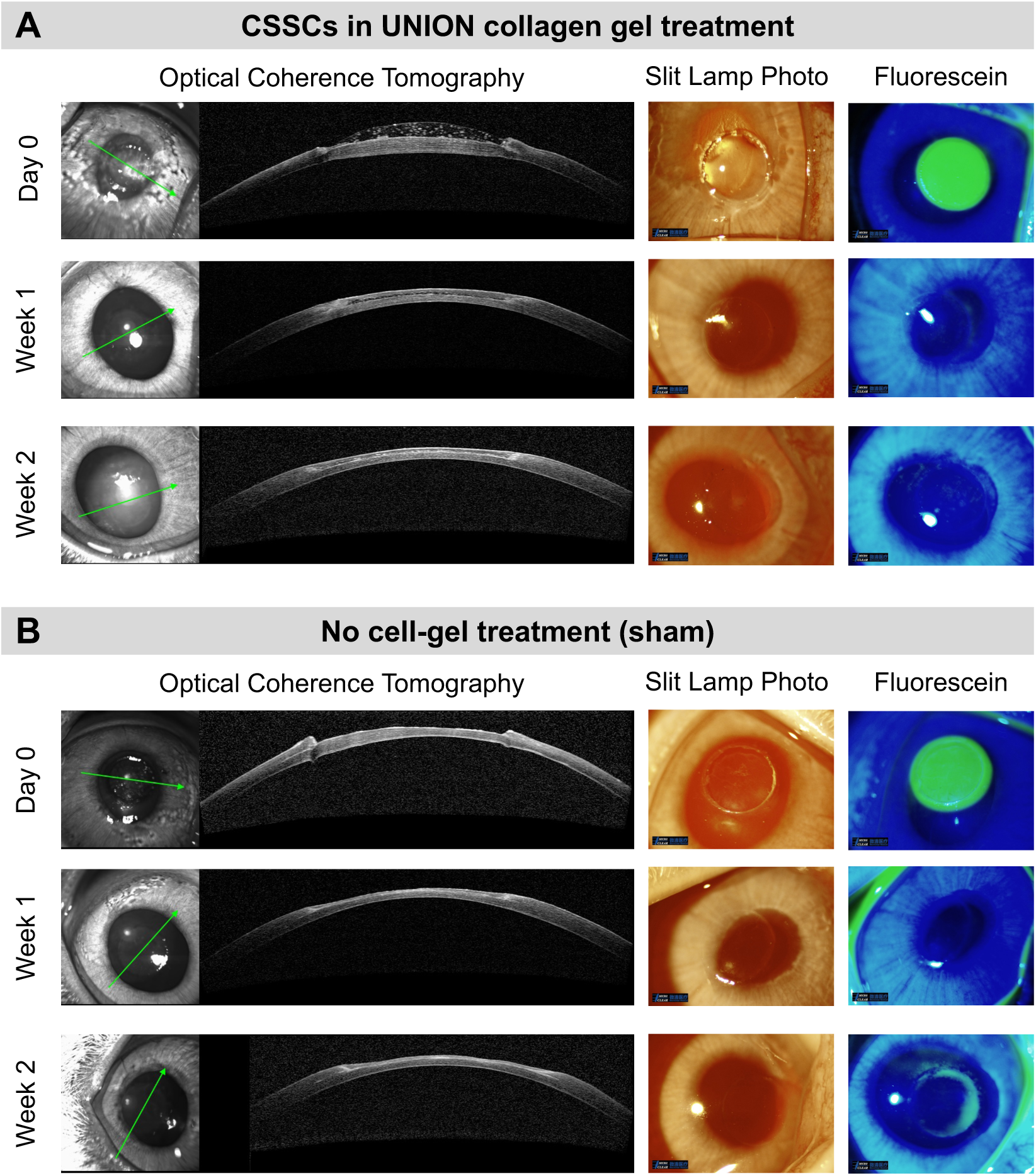
Clinical evaluation of corneal wounds in a rabbit anterior lamellar keratoplasty model. **(A)** Corneal wounds (3.5-mm diameter) were treated with CSSCs in UNION collagen that gelled *in situ*. Follow- up examinations with *in vivo* Optical Coherence Tomography (OCT), slit lamp photographs, and fluorescein wound staining were conducted on Week 1 and Week 2. N = 5 rabbits. **(B)** The sham controls did not receive the treatment of CSSCs in UNION collagen, but all other procedures for the surgery and treatment were kept constant. Follow-up examinations were similarly conducted on Week 1 and Week 2. N = 4 rabbits.

The corneas that received the cell-gel treatment (**Figure 7A**) and the sham control (**Figure 7B**) were clinically assessed immediately after the surgery (Day 0) with clinical follow-ups on Week 1 and Week 2. We used *in vivo* Optical Coherence Tomography (OCT) to visualize the corneal cross-section, slit lamp photographs to visualize the corneal transparency, and fluorescein staining under blue light to visualize the corneal wound (*i.e.,* where the corneal epithelium is missing). Immediately following the operation (Day 0), the UNION collagen gel restored the curvature of the cornea. In the clinical follow-ups (Week 1 and Week 2), the wounded corneas with the cell-gel therapy had a smooth transition between the keratectomy area and adjacent, uninjured corneal tissue. On the contrary, the sham control corneas without the cell-gel treatment had an open gap in the tissue on Day 0 where the keratectomy was performed, and a dip in the central cornea persisted over the 2 weeks. As expected, the removal of the corneal epithelium during the keratectomy resulted in a pronounced wound on Day 0 for both the cell-gel treated and sham corneas, as indicated by fluorescein staining. The corneas with the cell-gel treatment achieved complete re-epithelization within 1 week and did not experience any wound re-opening within 2 weeks. While the sham control group also re-epithelialized within 1 week, fluorescein staining after 2 weeks suggested that some wounds had re-opened. The treated corneas remained highly transparent, without the development of corneal haze. Furthermore, there was no evidence of inflammation, such as conjunctival infection, stromal infiltration, chamber flare, keratic precipitation, or corneal neovascularization.

To complement the *in vivo* evaluation in the rabbits, immunohistochemistry of the corneal stroma, epithelium, and endothelium indicates the regenerative effect of the cell-gel treatment (*i.e.*, CSSCs in UNION collagen bioink) for corneal wounds (**Figure 8, Figure S6**). After the Week 2 clinical evaluation, the rabbit eyes were enucleated and stained to assess the corneal tissue. The cell-gel treatment helped restore the thickness of the cornea in the wound center (**Figure S6A**) and the smooth transition between the wound and surrounding tissue at the wound edge (**Figure S6B**).

**Figure 8.**
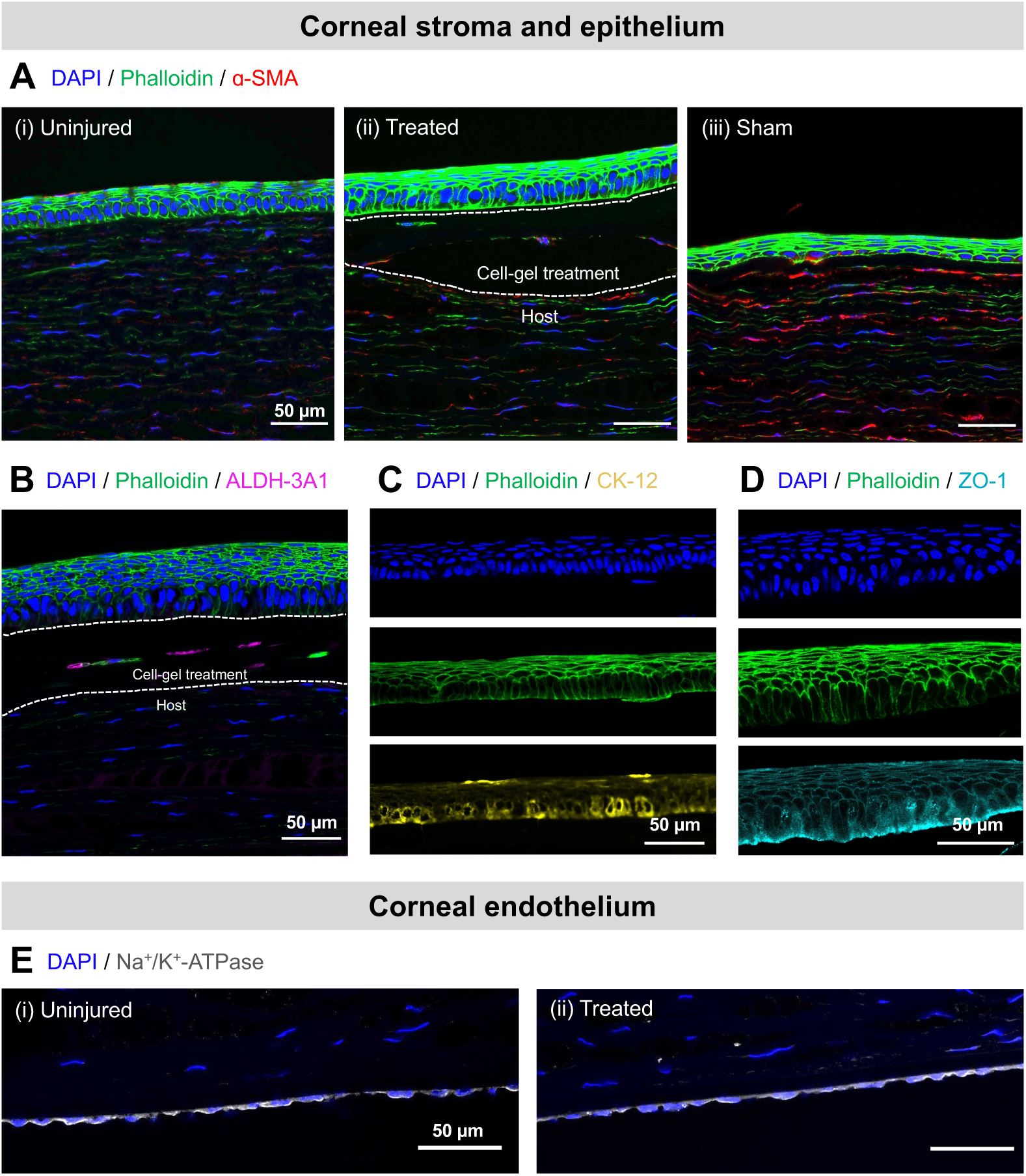
Immunohistochemical evaluation of the corneal stroma, epithelium, and endothelium in a rabbit anterior lamellar keratoplasty model (Week 2 post-operation). **(A)** Corneal stroma and epithelium of (i) uninjured corneas, (ii) treated corneas that received the cell-gel therapy, and (iii) control corneas that received a sham treatment after the keratectomy. Staining was performed with DAPI (nuclei, blue), phalloidin (F-actin, green), and anti-α-SMA (myofibroblast cell marker, red). Scale bars = 50 µm. **(B)** Corneal stroma and epithelium of treated corneas that received the cell-gel therapy. Staining was performed with DAPI (nuclei, blue), phalloidin (F-actin, green), and anti-ALDH-3A1 (human CSSC marker, magenta). **(C)** Corneal epithelium of treated corneas that received the cell-gel therapy. Staining was performed with DAPI (nuclei, blue), phalloidin (F-actin, green), and anti-CK-12 (corneal epithelial cell marker, yellow). **(D)** Corneal epithelium of treated corneas that received the cell-gel therapy. Staining was performed with DAPI (nuclei, blue), phalloidin (F-actin, green), and anti-ZO-1 (tight junctions, teal). **(E)** Corneal endothelium of (i) uninjured corneas and (ii) treated corneas that received the cell-gel therapy. Staining was performed with DAPI (nuclei, blue) and anti-Na^+^/K^+^-ATPase (endothelial pump marker, gray). Scale bars = 50 µm.

Keratocytes in the keratectomy area of the sham control corneal stroma were more activated than in the uninjured cornea or the cell-gel treated cornea, as demonstrated by immunofluorescence of alpha smooth muscle actin (α-SMA) (**Figure 8A**). The expression of α-SMA in keratocytes indicates the transformation from a quiescent phenotype into active myofibroblasts.(*49*) The presence and activity of myofibroblasts are known to correlate with undesired fibrotic responses and an increase in stromal haze.(*50*) Compared to the keratocytes in the corneas that had received the sham treatment, those in the corneas that received the cell-gel treatment expressed much less α-SMA and more closely resembled the uninjured stroma. Therefore, the cell-gel treatment helped avoid the myofibroblast transformation within the host stromal tissue that leads to corneal scarring. Furthermore, the human CSSCs delivered in the UNION collagen treatment expressed ALDH-3A1, the corneal crystallin characteristic of healthy CSSCs in the cornea (**Figure 8B**). The expression of ALDH-3A1 is lost during the transition of cells into myofibroblasts, so the continued expression of ALDH-3A1 by the transplanted CSSCs further confirms that they maintain their phenotype.(*49*, *51*) The anti-ALDH-3A1 antibody used is reactive with human cells but not rabbit cells, and therefore it does not stain rabbit cells from the host corneal tissue (**Figure S7**).

Additionally, differences were observed in the epithelium that regenerated over the surface of the corneal wounds (Figure 8A). The epithelium functions as a protective barrier, with a smooth surface and cell-to-cell tight junctions to prevent the entry of foreign materials.(*52*) Since no epithelial cells were seeded on top of the wounds as part of the treatment, the observed epithelium regenerated from the surrounding rabbit epithelium. The regenerated epithelium over the cell-gel treatment closely resembled the uninjured epithelium in its thickness and morphology, with columnar basal cells as the deepest epithelial layer and stratified squamous cells with flattened nuclei as the superficial layers (Figure 8A, panels i and ii). This tissue structure is characteristic of a healthy corneal epithelium.(*53*) In contrast, the regenerated epithelium in the sham control did not exhibit a similar morphology to the uninjured epithelium, with no layer of columnar cells present (Figure 8A, panel iii). The corneal epithelial cells over the cell-gel treatment also stained positively for CK-12 (**Figure 8C**), indicating their mature epithelial phenotype, and zonula occludens-1 (ZO-1) (**Figure 8D**), indicating the formation of tight junctions between cells in the multi-layered epithelium.

Finally, the corneal endothelium was examined for expression of the Na^+^/K^+^-dependent ATPase (**Figure 8E**). The Na^+^/K^+^-dependent ATPase is expressed in the basolateral membrane of the corneal endothelium and is responsible for its pump function.(*54*) After performing chemical reactions within the cornea, it is critical to confirm that there is not an adverse effect on the corneal endothelium. Corneal endothelial cells are known to be susceptible to damage and loss of pump function after common crosslinking techniques in the cornea, such as those requiring UV light.(*55*, *56*) In our approach, however, the bioorthogonal nature of the covalent click chemistry that crosslinks the UNION collagen does not create side-reactions or involve any cytotoxic molecules that could harm the surrounding, healthy tissue. Accordingly, the expression of Na^+^/K^+^-dependent ATPase in the endothelium after cell-gel treatment (Figure 8E, panel ii) was similar to the expression in the endothelium of the uninjured cornea (Figure 8E, panel i). Together, these immunohistochemical analyses reinforce the therapeutic potential of the UNION collagen bioink to deliver CSSCs and promote corneal regeneration, by effectively minimizing fibrotic responses and restoring the integrity of the stromal and epithelial layers without compromising the corneal endothelium.

## DISCUSSION

The ability to reconstruct and regenerate ulcerated and scarred corneas on demand would be of great benefit to patients suffering from corneal blindness. Together, our *in vitro* and *in vivo* results demonstrate the promising therapeutic properties of UNION collagen bioinks that deliver CSSCs directly to corneal wounds with a bioorthogonal, *in situ* gelation mechanism. The covalent crosslinking chemistry allows the cell-gel therapy to be optically transparent, stable against contraction forces exerted by CSSCs, and permissive to the efficient growth of corneal epithelial cells. CSSCs remain alive within the UNION collagen gel precursor solution under standard storage and transportation conditions, and their clinical potential to promote corneal wound healing and re-epithelialization was demonstrated in a rabbit anterior lamellar keratoplasty model.

Designing laboratory-made constructs with the long-term transparency and regenerative capacity of a human donor cornea has been a formidable challenge in the field of ophthalmic tissue engineering. The off-label use of cyanoacrylate glue is commonly used to stabilize acute corneal perforations.(*57*) However, the glue creates an opaque, rough surface, and further interventions such as emergency corneal transplantations are often required.(*58*, *59*) Previous biomaterial-based strategies have commonly included pre-formed, implanted buttons composed of various acellular matrices such as collagen, gelatin, and hyaluronic acid,(*15*, *60–63*) as well as acellular *in situ*-forming matrices that are photocrosslinked or chemically crosslinked within corneal wound sites.(*12*, *13*, *22*, *64*, *65*) Acellular biomaterials offer structural support to the cornea but do not provide the cellular components that are present in corneal grafts. Therefore, there is growing interest in stem cell therapies for corneal regeneration.(*66*) This shift is underscored by the European Commission’s recent approval of Holoclar—a limbal stem-cell based product to replace the corneal epithelium after an ocular burn—for use throughout the European Union.(*67*)

In our work, we aimed to advance corneal regeneration by targeting not only the replacement of the corneal epithelium, but also the regeneration of the corneal stroma after an injury. CSSCs are an attractive therapeutic avenue, since they can be obtained by biopsy and are available as clinical-grade cultures from human cadaveric corneas,(*68*, *69*) providing a ready source of expandable stem cells.(*9*, *10*) The reproducible derivation and expansion of CSSCs has been well-established, with a potential yield from each cornea of 12 to 16 x 10^10^ CSSCs upon five passages.(*5*) Therefore, a single donor cornea could supply enough CSSCs to treat thousands of patients with the UNION collagen bioink therapy, a significant advantage over traditional corneal transplants that require a 1:1 donor cornea-to-patient ratio. In this study, we delivered CSSCs directly to corneal wounds within an engineered UNION collagen matrix, providing both structural and cellular support for effective tissue regeneration. Importantly for cellular therapies, we demonstrated that CSSCs survive cold storage conditions within a UNION collagen precursor solution, which would be required for the transportation and distribution of the therapy to surgery centers. Our results indicate that this strategy could integrate into existing infrastructures for the transit of donor corneas. Additionally, the *in situ* gelation mechanism of the UNION collagen bioink makes it a targeted therapy that replaces only the damaged corneal tissue while preserving surrounding healthy tissue. This approach minimizes the need for invasive surgeries and full corneal replacements using donor tissue, which is severely limited in supply and availability worldwide. The use of UNION collagen bioinks to directly deliver CSSCs to corneal wounds may therefore serve as a practically and clinically feasible solution to address the dearth of corneal graft tissue available for transplantation.

Looking ahead, our findings open several opportunities for future investigation. Given the high viability of CSSCs after cold storage under typical shipping conditions, more challenging storage scenarios—such as extended time or temperature fluctuations—could be studied to determine the limits of the CSSCs’ distribution, particularly for under-resourced areas. The promising pre- clinical results at the Week 2 timepoint motivate longer *in vivo* studies to confirm long-term corneal regeneration and to study the degradation or persistence of the UNION collagen gel and transplanted CSSCs over time. To identify the range of corneal wound sizes for which this cell-gel therapy is effective, wounds with greater diameters than 3.5 mm and deeper cuts than 50% of the corneal thickness could be assessed. The *in situ* gelling bioink described here may be less suited for restoring the corneal curvature of wider wounds. *Ex situ* bioprinting of the UNION collagen could therefore be used to achieve the patient-specific shape, size, and curvature of the engineered corneal graft. Since the UNION bioprinting approach is amenable to multi-material constructs with cohesive interfaces,(*25*) it could be adapted to include other extracellular matrix components in the UNION bioink in addition to collagen, such as hyaluronic acid,(*65*) or to create multi-layered corneal substitutes with patterning of different matrix and cellular compositions for full-scale corneal implants.

Overall, this work represents an important step toward the development of a regenerative medicine therapy for patients suffering from corneal blindness. By leveraging the unique properties of UNION collagen bioinks to directly deliver regenerative CSSCs to corneal wounds, this approach has the potential to serve as an effective alternative to transplantation of cadaveric human corneas.

## MATERIALS AND METHODS

### Preparation of UNION collagen gels

Collagen-azide was synthesized by modifying type I bovine atelocollagen solution (5 mg/mL, Gibco) with azide functional groups using N-hydroxysuccinimide (NHS) ester chemistry to react with primary amines on collagen.(*26*, *64*) First, the acidic collagen solution was neutralized on ice following instructions from the manufacturer, using 1.0 M sodium hydroxide solution (NaOH, Sigma), ultrapure deionized water (Millipore), and 10X phosphate buffered saline (PBS, Millipore) to reach a pH of 7.5 and a concentration of 4 mg/mL collagen. Azido-PEG4-NHS ester (BroadPharm) was dissolved in dimethyl sulfoxide (DMSO, Fisher) at a concentration of 100 mg/mL and added to the neutralized collagen solution at 2 molar equivalents relative to primary amines on the collagen. The solution was mixed well, rotated for 2 h at 4 °C, and then dialyzed overnight in a Slide-A-Lyzer dialysis kit (3.5-kDa MWCO, ThermoScientific) against 1X PBS at 4 °C. The purified collagen-azide solution was stored at 4 °C before use. The degree of functionalization was determined using a 2,4,6-Trinitrobenzene Sulfonic Acid (TNBSA) assay (Thermo Scientific) to quantify free amino groups, following the instructions from the manufacturer.

The PEG-BCN crosslinker was synthesized as previously described.(*25*) In brief, PEG-amine (4 arm, 20-kDa, Creative PEGworks) was dissolved at 10 mg/mL in anhydrous DMSO. Then, (1*R*, 8*S*, 9*S*)-bicyclo[6.1.0]-non-4-yn-9ylmethyl *N*-succinimidyl carbonate (BCN-NHS, 1 molar equivalent relative to amines, Sigma) and triethylamine (1.5 molar equivalent relative to amines, Fisher) were added dropwise. The reaction was purged with nitrogen gas and proceeded overnight at room temperature with constant stirring. The solution was then dialyzed against ultrapure deionized water for 3 days, sterile filtered through a 0.22 µm filter, lyophilized, and stored at -80 °C before use.

For the preparation of UNION collagen hydrogels, lyophilized PEG-BCN crosslinker was dissolved in PBS (acellular studies) or CSSC cell culture medium (cellular studies) and then mixed with the collagen-azide to reach a final concentration of 3 mg/mL collagen-azide and 4 mg/mL PEG-BCN, unless specified otherwise. After thorough mixing, the solution was pipetted into molds and incubated at 37 °C for crosslinking and gelation.

### Preparation of PHYS collagen gels

For the preparation of PHYS collagen gels, type I bovine atelocollagen solution (either 5 mg/mL, Gibco; or 10 mg/mL, Advanced BioMatrix) was neutralized on ice following instructions from the manufacturers, using 1.0 M NaOH (Sigma), ultrapure deionized water (Millipore), and 10X PBS (Millipore) to reach a pH of 7.5. The initial 5 mg/mL and 10 mg/mL collagen solutions were used to prepare PHYS collagen gels with concentrations less than 5 mg/mL and between 5-10 mg/mL, respectively. Once neutralized and diluted with PBS (acellular studies) or CSSC cell culture medium (cellular studies) to reach the appropriate concentration, the solution was pipetted into molds and incubated at 37 °C for gelation.

### Hydrogel characterization

Oscillatory shear measurements were conducted on an ARG2 rheometer (TA Instruments) equipped with a Peltier plate and a solvent trap to prevent evaporation. A cone-plate geometry with a 1° angle and 20-mm diameter was used. Time-sweep measurements to study the gelation kinetics were carried out at an angular frequency of 1 rad/s and a shear strain amplitude of 1%. Frequency-sweep measurements (between 10^-1^ and 10^2^ rad/s) were conducted at a shear strain amplitude of 1%. All measurements were confirmed to be within the linear viscoelastic regime.

The transmittance of the hydrogels was calculated based on the measured absorbance. First, 50 μL of gel were prepared within the wells of clear 96-well plates. The absorbance was measured between 300 and 800 nm (SpectraMax M2 Microplate Reader, Molecular Devices) with PBS as a blank. The transmittance was calculated using the relationship T (%) = 1/10^A^ × 100, where A is the absorbance.

For visualization of fibrillar microstructures within gels, second harmonic generation (SHG) imaging was conducted using an inverted microscope (Nikon, Ti2-E equipped with a C2 confocal scanning head and a Nikon CFI Plan Apo IR 60XC water immersion objective). The SHG signal was generated by probing the samples with a picosecond-pulsed laser from a system (APE America Inc., picoEmerald S with 2 ps pulse length, 80 MHz repetition rate, and 10 cm^-1^ bandwidth) consisting of an optical parametric oscillator (OPO) tunable between 700-960 nm and pumped by the SHG of a 1031 nm mode-locked ytterbium fiber laser. The OPO wavelength was set to 797 nm. The backscattered SHG signal (at a wavelength of 398.5 nm) was separated using a set of optical filters (BrightLine 400/12 bandpass, BrightLine 390/18 bandpass, Thorlabs FESH0500 shortpass) and detected pixel-by-pixel with a photomultiplier tube (Hamamatsu, R6357), with a pixel size of <80 nm and dwell time of 10.8 s. The excitation power at the sample was 50 mW.

Hydrogel contraction imaging was performed with an epifluorescent microscope (Leica Microsystems, THUNDER Imager 3D Cell Culture) with a 2.5X air objective in bright field mode. To track the contraction of the hydrogels over 7 days, tile-scan images of the gels within silicone molds (8-mm diameter) bonded to glass coverslips were taken at each time point, and their areas were measured with the open-source image analysis software FIJI (ImageJ2, Version 2.3.0/1.53f).(*70*)

### CSSC culture and encapsulation in gels

The CSSCs were isolated from human corneas (Lions Eye Institute for Transplant and Research) as previously described.(*5*) Briefly, the endothelial layer was first removed, followed by the central corneal button using an 8-mm trephine. The remaining cornea-scleral rims were cut into three pieces and placed epithelial-side down on tissue culture plastic in a 6-well plate. The segments attached to the tissue culture plastic after air-drying for 2 min. CSSC growth medium was prepared with 500 mL MEM-α (Corning), 50 mL fetal bovine serum (Gibco), 5 mL GlutaMax (Gibco), 5 mL non-essential amino acids (Gibco), and 5 mL antibiotic-antimycotic (Gibco). Drops of the CSSC growth medium were added to the cornea-scleral rim segments twice each day until the cells formed a matrix that was fully attached to the tissue culture plastic. After that point, 1-2 mL of CSSC growth medium were added every other day. Once the cells reached 50% confluency, the cornea-scleral rim segments were removed, and the cells in the well were trypsinized and plated into tissue culture flasks (Thermo Fisher Scientific). Growth medium was changed every other day, and CSSCs were passaged upon reaching 80% confluency. For cell encapsulations, the CSSCs were trypsinized, counted, pelleted, and re-suspended at a density of 3 x 10^6^ cells/mL within the UNION collagen bioink solution (3 mg/mL collagen-azide, 4 mg/mL PEG-BCN). The solution was then pipetted into silicone molds bonded to glass coverslips and allowed to gel for 1 h at 37 °C. The growth medium was changed every other day during the duration of the culture period.

### Corneal epithelial cell culture and seeding on gels

Human telomerase corneal epithelial cells (CECs) were kindly donated by Dr. Ali Djalilian (University of Illinois at Chicago, USA). CEC growth medium was prepared with 500 mL keratinocyte SFM cell medium (Gibco), 5 mL penicillin-streptomycin (Sigma-Aldrich), 2.5 mL hydrocortisone (STEMCELL Technologies), and 250 µL insulin (Sigma-Aldrich). Before plating the cells, flasks were coated with 3 mL of FNC Coating Mix (Athena Enzyme Systems) for 3 min at room temperature. Growth medium was changed every other day, and CECs were passaged upon reaching 80% confluency. For seeding atop hydrogels, the CECs were trypsinized, counted, pelleted, re-suspended in media, and seeded at an initial cell density of 25,000 cells/cm^2^. When the gels also contained encapsulated CSSCs, a 1:1 ratio of the CSSC and CEC growth media was used. The cell culture medium was changed every other day during the duration of the culture period.

### *In vitro* cell characterization

To assess the viability of the CSSCs, a Live/Dead assay was conducted by staining cells with calcein AM and ethidium homodimer-1 (Life Technologies), following the manufacturer’s instructions. Imaging was performed using a STELLARIS 5 confocal microscope (Leica) with a 10X air objective. At least 3 images were taken in different areas of each sample, and image analysis was performed using FIJI (ImageJ2, Version 2.3.0/1.53f). Cell viability was calculated as the number of live cells (calcein AM-positive) divided by the total number of cells.

The secretome of CSSCs growing on 2D tissue culture plastic or encapsulated in PHYS or UNION collagen gels was analyzed with a human cytokine/chemokine/growth factor panel A bead-based multiplex panel, using the Luminex xMAP technology (Millipore Sigma). The Luminex assay uses a mixture of color-coded beads coated with analyte-specific antibodies, and the captured analytes are detected with biotinylated detection antibodies. The secretome samples were collected on Day 5 after CSSC encapsulation, frozen at -80 °C, and transferred to the Stanford Human Immune Monitoring Center (HIMC) on dry ice for analysis, avoiding freeze-thaw cycles. The expression of analytes for the secretome from CSSCs in PHYS or UNION collagen gels was normalized to the secretome of the same number of CSSCs growing on 2D tissue culture plastic.

For immunofluorescence imaging, gels with encapsulated CSSCs or seeded CECs were fixed with 4% paraformaldehyde (PFA) in PBS for 30 min at room temperature (RT). Samples were permeabilized for 1 hour at RT with 0.25% Triton X-100 (Sigma Aldrich) in PBS (PBST) and then blocked for 3 hours at RT in PBS with 5 wt% bovine serum albumin (BSA, Roche), 5% goat serum (Gibco), and 0.5% Triton X-100. Antibody dilutions were prepared in PBS with 2.5 wt% BSA, 2.5% goat serum, and 0.5% Triton X-100. Each sample was first treated with the appropriate primary antibody overnight at 4 °C: for CSSCs, rabbit anti-ALDH-3A1 (Abcam, ab76976, 1:200 dilution); for CECs, mouse Alexa Fluor 488 anti-Ki-67 (Abcam, ab281847, 1:100 dilution) and rabbit Alexa Fluor 488 anti-CK-12 (Abcam, ab222116, 1:100 dilution). For staining of hyaluronic acid, biotinylated hyaluronan binding protein (Millipore Sigma, 1:100 dilution) was used in place of a primary antibody. The following day, the samples were washed with PBST and incubated with a corresponding fluorescently tagged secondary antibody overnight at 4 °C, if the primary antibody did not already contain a fluorophore. For staining of hyaluronic acid, Alexa Fluor 488 Streptavidin (Invitrogen, 1:500 dilution) was used in place of a secondary antibody. The following day, samples were washed again with PBST and incubated with 4′,6-diamidino-2-phenylindole (DAPI, Molecular Probes, 1:2000 dilution) and phalloidin-tetramethyl rhodamine B isothiocyanate (phalloidin-TRITC, Sigma Aldrich, 1:100 dilution) in PBST for 1 h at RT. Finally, samples were washed with PBST, mounted onto glass slides with ProLong Gold Antifade Mountant (Thermo Fisher Scientific), and allowed to cure for 48 hours before imaging with a STELLARIS 5 confocal microscope (Leica). Image analysis of the confluence and proliferation of CECs on the gels was conducted with CellProfiler, an open-source software for measuring and analyzing cell images.(*71*)

### *In vivo* rabbit anterior lamellar keratoplasty model and evaluation

Animal experiments were designed to conform with the Association for Research in Vision and Ophthalmology (ARVO) Statement for the Use of Animals in Ophthalmic and Vision Research, and the protocols were reviewed and approved by the Stanford University Institutional Animal Care and Use Committee (IACUC). Adult New Zealand White rabbits (3-4 kg) were used for this study. All anesthesia procedures were performed by the Veterinary Service Center (VSC) staff at Stanford University. An anterior lamellar keratectomy was performed on the rabbit’s right eye using a 3.5-mm trephine to create a circular cut and a spatula to remove the tissue. Before applying any treatment, the cornea was imaged with optical coherence tomography (OCT, Heidelberg Engineering) to confirm that the depth of the wound was ∼50% of the corneal thickness. Then, 5-7 μL of CSSCs within the collagen-azide and PEG-BCN solution were applied with a pipette to fully fill the wound site for the treated group, and saline was applied for the sham group. The cell-gel therapy was allowed to gel *in situ* for 10 min before initial evaluation.

Corneas were examined *in vivo* with OCT images and photographs. The lack of an epithelium at the wound site was identified using fluorescein solution (1.5 wt%) that was dropped onto the eye and then washed immediately with balanced salt solution (BSS, Alcon). The fluorescein-stained areas were photographed under cobalt blue light. A contact lens was applied to protect the gel from scratching by the rabbit, and a temporary tarsorrhaphy was performed to prevent agitation of the rabbit and to help keep the contact lens and gel in place for the first 7 days. Ofloxacin and 1% prednisolone acetate ophthalmic eye drops were applied 3 times daily to prevent infection and to retain moisture of the eye. Follow-up examinations were conducted on Days 7 and 14.

To obtain tissues for analysis, rabbits were euthanized on Day 14 according to protocols consistent with the recommendations of the Panel of Euthanasia of the American Veterinary Medical Association. After enucleation, corneas were fixed in 4% PFA, embedded in optimal cutting temperature compound, and cryosectioned in preparation for immunohistochemistry. The corneal sections were treated with the following primary antibodies: mouse anti-α-SMA (Abcam, ab7817, 1:100 dilution), rabbit anti-ALDH-3A1 (Abcam, ab76976, 1:200 dilution), rabbit Alexa Fluor 488 anti-CK-12 (Abcam, ab222116, 1:100 dilution), rat anti-ZO-1 (Thermo Fisher Scientific, 14-9776-82, 1:50 dilution), and mouse anti-Na^+^/K^+^-ATPase (Abcam, ab7671, 1:200 dilution). If the primary antibody did not already contain a fluorophore, corneal sections were incubated with a corresponding fluorescently tagged secondary antibody overnight at 4 °C. Additionally, nuclei were stained with DAPI (Invitrogen, 1:2000 dilution), and F-actin was stained with Alexa Fluor 488 Phalloidin (Molecular Probes, 1:40 dilution). Stained corneal tissue sections were imaged with a STELLARIS 5 confocal microscope (Leica).

### Statistical analysis

Statistical analyses were performed using GraphPad Prism (Version 10). Details of the sample sizes and statistical tests conducted for each figure are included within the figure captions. In all cases, statistical differences are denoted as follows: not significant (ns, *p* > 0.05), * (*p* < 0.05), ** (*p* < 0.01), *** (*p* < 0.001), and **** (*p* < 0.0001).

## ACKNOWLEDGEMENTS

The authors thank Annika Enejder for microscope use for second harmonic generation imaging; the veterinary technicians at the Stanford School of Medicine Veterinary Service Center for their assistance caring for the rabbits used in the *in vivo* studies; Roopa Dalal for assistance fixing and sectioning corneas of enucleated rabbit corneas; and the Stanford Human Immune Monitoring Center (HIMC), especially Yael Rosenberg-Hasson and Holden Maecker, for assistance with the cytokine immunoassay. Schematics included within Figures 6, S3, and S4 were created with Biorender.com.

## Author contributions

Conceptualization: L.G.B., S.C.H., D.M. Methodology: L.G.B., B.C., S.M.H., S.C.H., D.M. Investigation: L.G.B., B.C., S.M.H., U.H., T.W., G.F-C., Y.A.S., P.K.J. Visualization: L.G.B., B.C., S.M.H., U.H., T.W. Funding acquisition: S.C.H., D.M. Supervision: S.C.H., D.M. Writing – original draft: L.G.B., S.C.H., D.M. Writing – review and editing: L.G.B., B.C., S.M.H., U.H., T.W., G.F-C., Y.A.S., P.K.J., S.C.H., D.M.

## Data and materials availability

All data needed to evaluate the conclusions in the paper are present in the paper and/or Supplementary Materials.

## LIST OF SUPPLEMENTARY MATERIALS

Figs. S1 to S7

