## Supplementary Material for "In Situ UNIversal Orthogonal Network (UNION) Bioink Deposition for Direct Delivery of Corneal Stromal Stem Cells to Corneal Wounds"

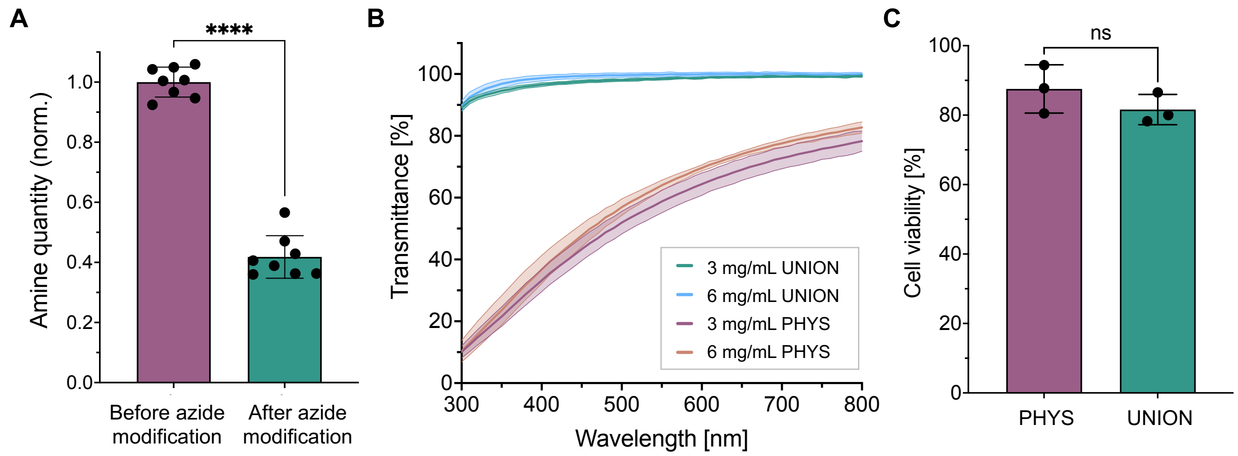


**Figure S1. UNION collagen properties. (A)** Approximately 40% of the primary amines in collagen remain after the collagen-azide bioconjugation reaction, indicating that approximately 60% of the primary amines are modified with azides. Normality of the data was confirmed with the Shapiro-Wilk test, and statistical analysis was performed with an unpaired t test. N = 8 independent samples per material condition. Data plotted as mean ± SD. ****$p$ < 0.0001. **(B)** UNION collagen gels have greater transmittance of light than PHYS collagen gels across the visible light regime for both 3 mg/mL and 6 mg/mL collagen conditions. N = 3 independent gels per material condition. Shaded regions represent the standard deviation from the mean. **(C)** The viability of encapsulated CSSCs remains similar for PHYS and UNION collagen after 14 days in culture. Normality of the data was confirmed with the Shapiro-Wilk test, and statistical analysis was performed with an unpaired t test. N = 3 independent samples per material condition. Data plotted as mean ± SD. ns = not significant.

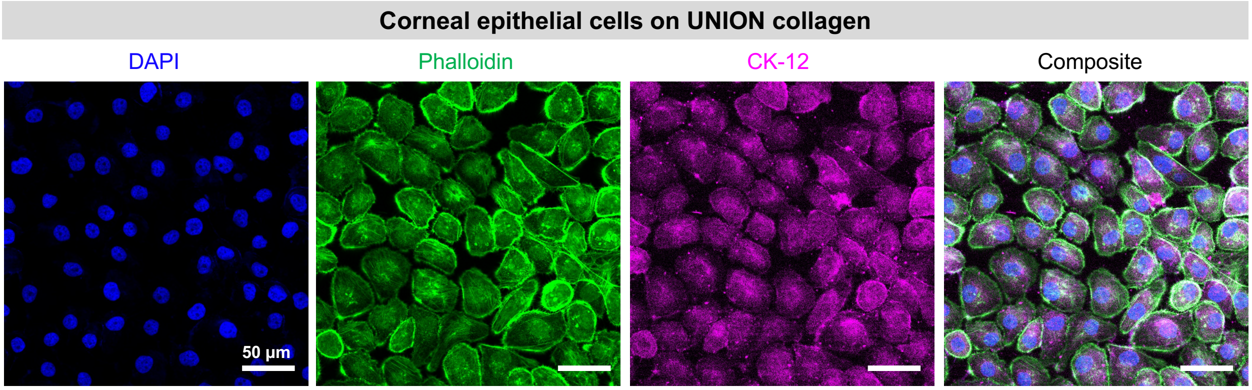


**Figure S2. Corneal epithelial cells express cytokeratin 12 (CK-12) while growing on UNION collagen gels.** Representative images of staining with DAPI (nuclei, blue), phalloidin (F-actin, green), and anti-CK-12 (corneal epithelial cell marker, magenta). Scale bars = 50 μm.

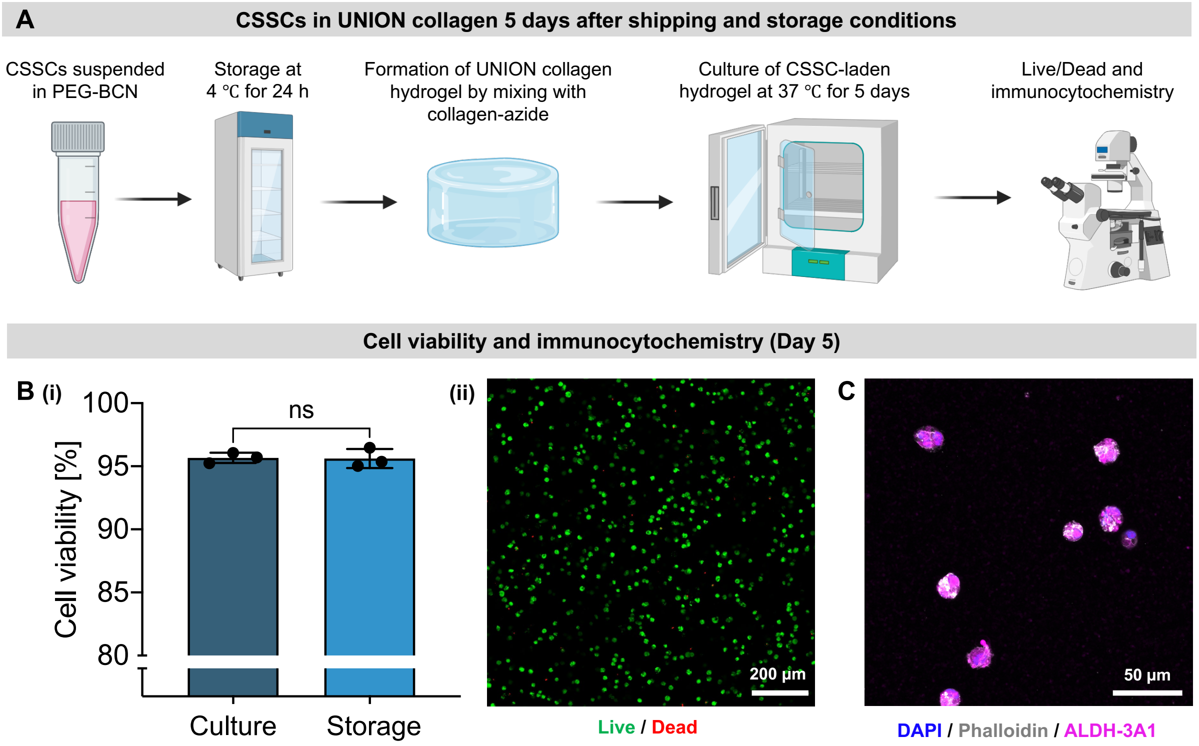


**Figure S3. CSSCs cultured in UNION collagen for 5 days after exposure to cold storage conditions for 24 h. (A)** Schematic illustrating the procedure to evaluate CSSCs in UNION collagen 5 days after their cold storage. CSSCs are suspended in PEG-BCN in cell culture medium and stored for 24 h at 4 °C. A CSSC-laden UNION collagen gel is then formed by mixing with collagen-azide and then cultured at physiological temperature (37 °C) for 5 days. **(B)** (i) Quantification of CSSC viability in UNION collagen 5 days after cell encapsulation, comparing cells that had been prepared with normal cell culture conditions or the cold storage conditions. N = 3 independent samples per condition. Normality of the data was confirmed with the Shapiro-Wilk test, and statistical analysis was performed with an unpaired t test. Data plotted as mean ± SD. ns = not significant. (ii) Representative image from Live / Dead cytotoxicity assay. **(C)** Representative image of staining with DAPI (nuclei, blue), phalloidin (F-actin, gray), and anti-ALDH-3A1 (CSSC marker, magenta).

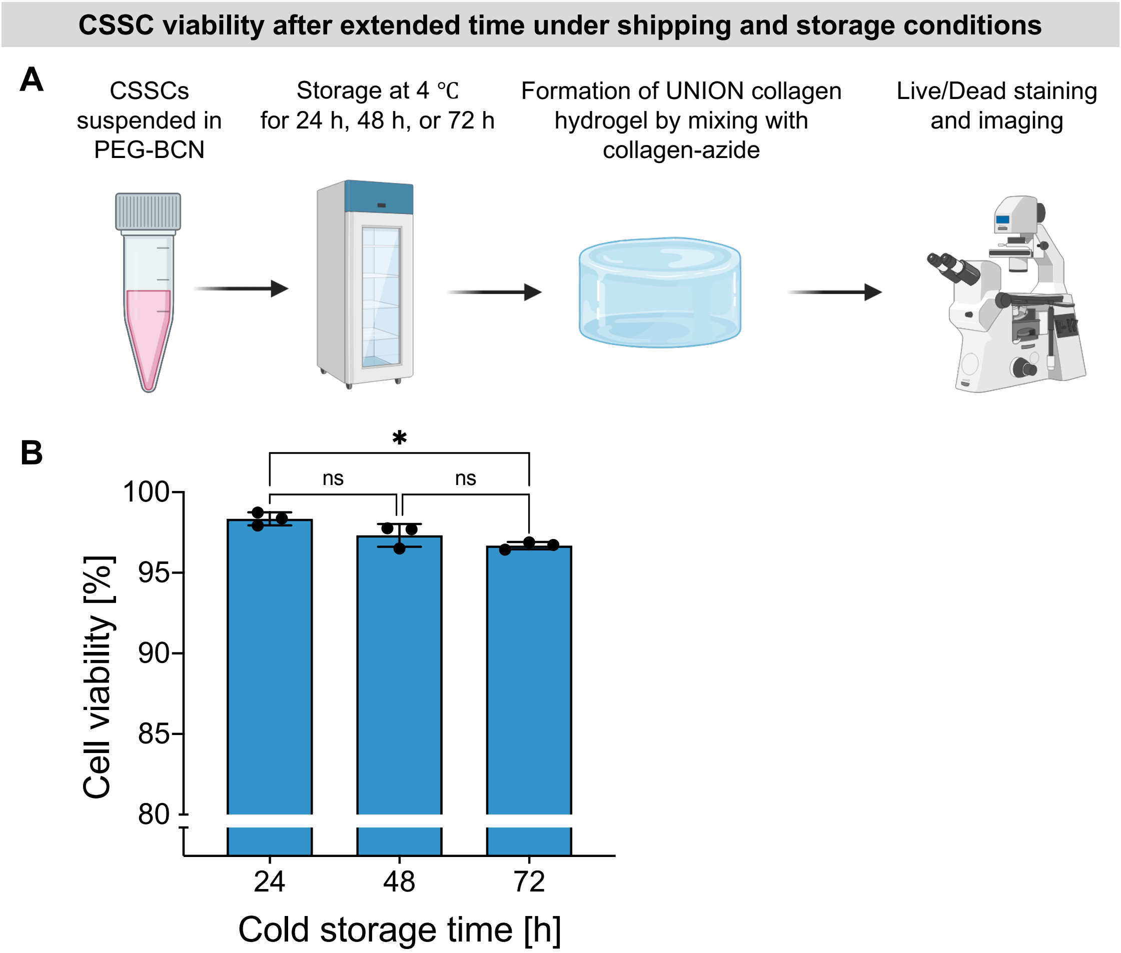


**Figure S4. CSSC viability after extended cold storage conditions. (A)** Schematic illustrating the procedure to evaluate CSSCs after extended times in cold storage. CSSCs are suspended in PEG-BCN in cell culture medium and stored at 4 °C for 24 h, 48 h, or 72 h. A CSSC-laden UNION collagen gel is then formed by mixing with collagen-azide, and the cell viability is assessed with a Live / Dead cytotoxicity assay. **(B)** Quantification of CSSC viability in UNION collagen to compare cells that had been in cold storage at 4 °C for 24 h, 48 h, or 72 h. N = 3 independent samples per condition. Normality of the data was confirmed with the Shapiro-Wilk test, and statistical analysis was performed with an unpaired t test. Data plotted as mean ± SD. ns = not significant. *$p < 0.05$.

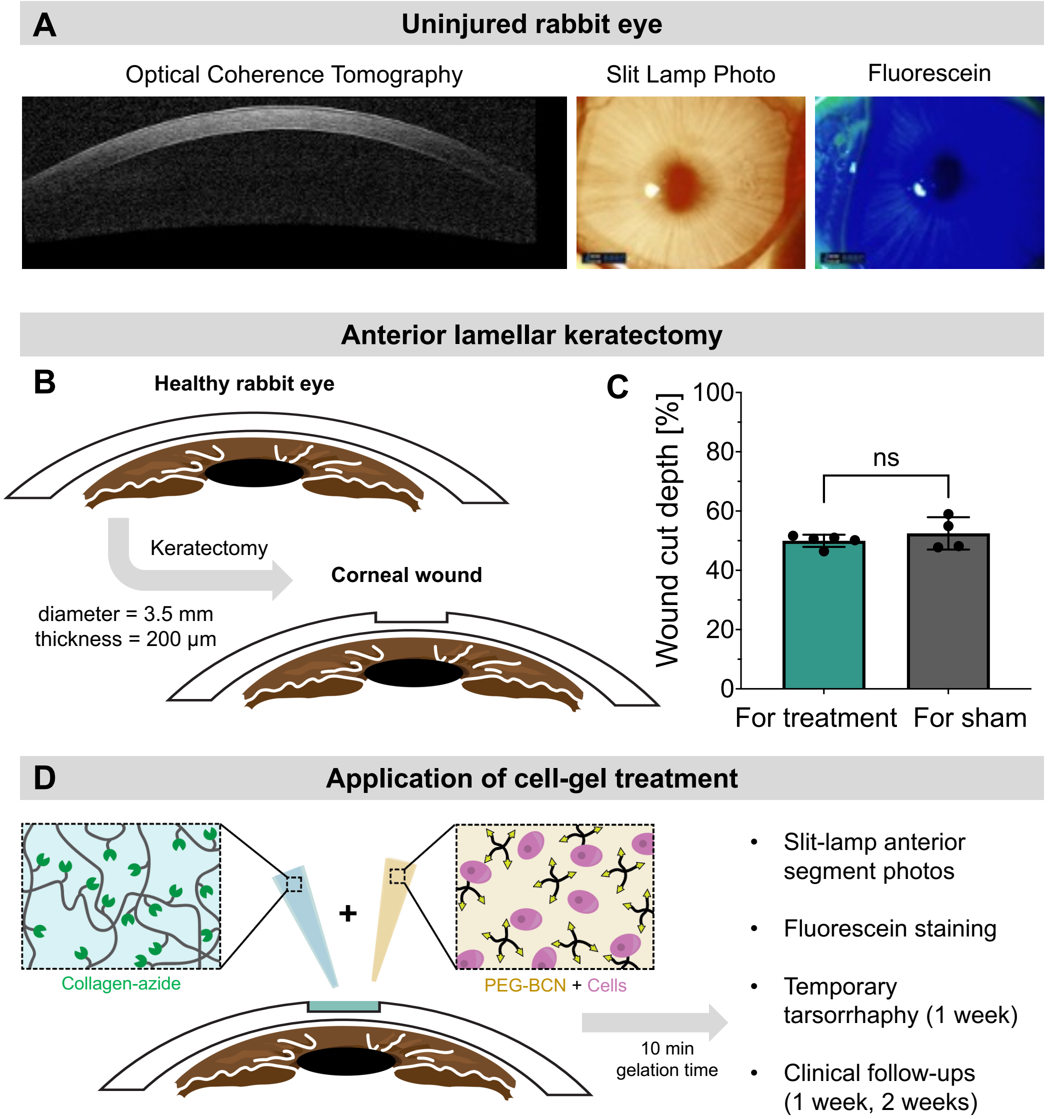


**Figure S5. Rabbit anterior lamellar keratoplasty model. (A)** *In vivo* Optical Coherence Tomography (OCT), slit lamp photographs, and fluorescein wound staining of uninjured corneas confirm that there are no corneal abnormalities prior to the keratectomy. **(B)** A keratectomy on the rabbit eye creates a corneal wound with a diameter of 3.5 mm and thickness of 200 µm. **(C)** The depth of the wound is controlled to be approximately 50% of the thickness of the rabbit cornea. Normality of the data was confirmed with the Shapiro-Wilk test, and statistical analysis was performed with an unpaired t test. N = 5 rabbits for the cell-gel treatment. N = 4 rabbits for the sham treatment control group. Data plotted as mean ± SD. ns = not significant. **(D)** To apply the cell-gel treatment of CSSCs in UNION collagen, the precursor solutions (collagen-azide; PEG-BCN with CSSCs) are first mixed together to create the bioink. The bioink is then deposited into the corneal wound to fill the defect site and allowed to gel for 10 mins before clinical evaluation.

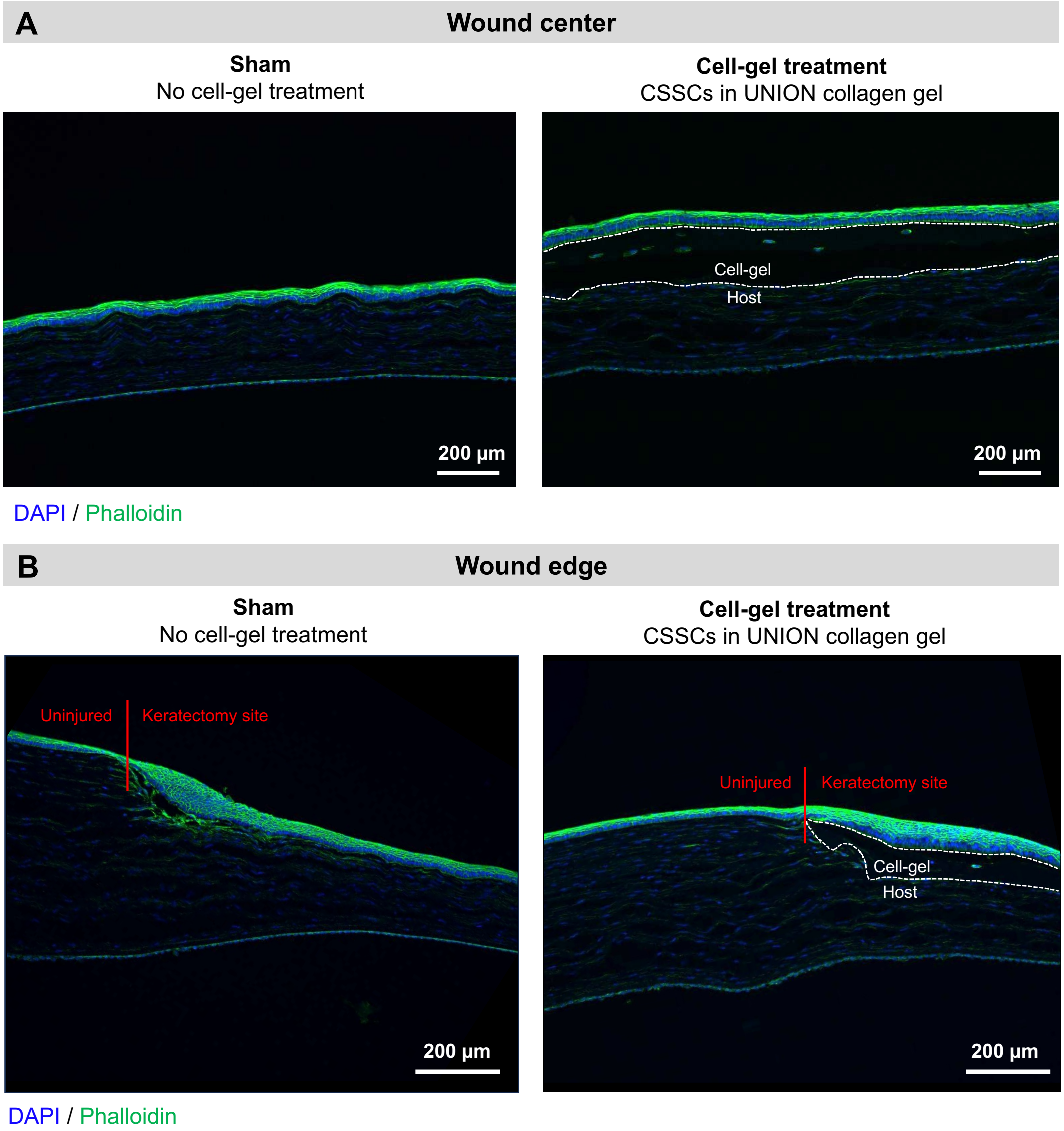


**Figure S6. Corneal tissue after enucleation during Week 2 post-operation. (A)** Comparison of wound center between sham treatment (left) and cell-gel treatment (right). **(B)** Comparison of wound edge between sham treatment (left) and cell-gel treatment (right). All staining was performed with DAPI (nuclei, blue) and phalloidin (F-actin, green).

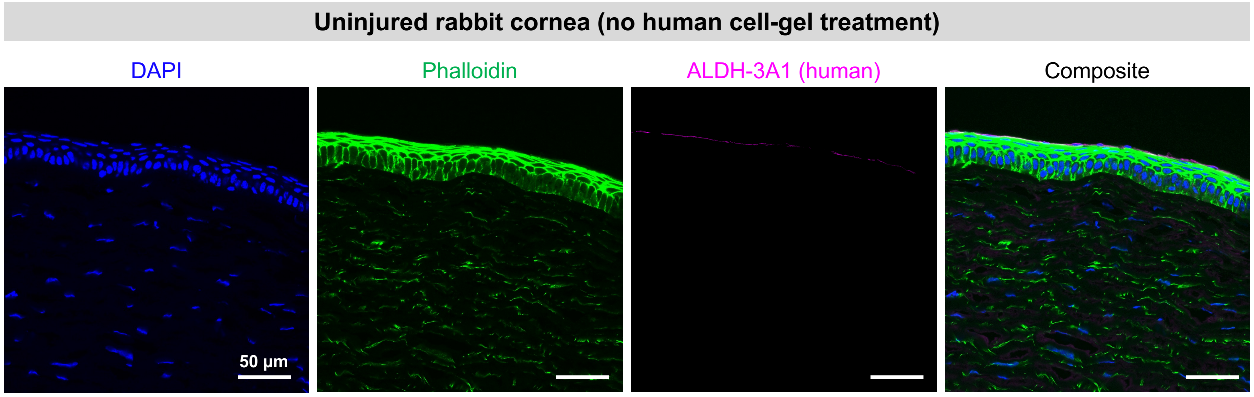


**Figure S7. Cells in uninjured rabbit corneas do not stain positive for human ALDH-3A1.** Representative images of staining with DAPI (nuclei, blue), phalloidin (F-actin, green), and anti-ALDH-3A1 (human CSSC marker, magenta). The anti-ALDH-3A1 antibody used is reactive with human cells but not rabbit cells, and therefore it does not stain rabbit cells from the host corneal tissue. Scale bars = 50 µm.
